# High-throughput mapping of long-range neuronal projection using *in situ* sequencing

**DOI:** 10.1101/294637

**Authors:** Xiaoyin Chen, Yu-Chi Sun, Huiqing Zhan, Justus M Kebschull, Stephan Fischer, Katherine Matho, Z. Josh Huang, Jesse Gillis, Anthony M Zador

## Abstract

Understanding neural circuits requires deciphering interactions among myriad cell types defined by spatial organization, connectivity, gene expression, and other properties. Resolving these cell types requires both single neuron resolution and high throughput, a challenging combination with conventional methods. Here we introduce BARseq, a multiplexed method based on RNA barcoding for mapping projections of thousands of spatially resolved neurons in a single brain, and relating those projections to other properties such as gene or Cre expression. Mapping the projections to 11 areas of 3579 neurons in mouse auditory cortex using BARseq confirmed the laminar organization of the three top classes (IT, PT-like and CT) of projection neurons. In depth analysis uncovered a novel projection type restricted almost exclusively to transcriptionally-defined subtypes of IT neurons. By bridging anatomical and transcriptomic approaches at cellular resolution with high throughput, BARseq can potentially uncover the organizing principles underlying the structure and formation of neural circuits.

## Introduction

The nervous system consists of networks of neurons, organized into areas, nuclei, lattices, laminae and other structures. Within these structures are uncounted different neuronal types, each characterized by its own pattern of connections, gene expression and physiological properties. Understanding how diverse neurons are organized thus requires methods that can map various neuronal characteristics at cellular resolution, with high-throughput, in single brains.

A particular challenge is to map the long-range axonal projection patterns that form the physical basis for neuronal circuits across brain areas. Traditional neuroanatomical methods based on microscopy can be used to visualize neuronal morphology, including long-range projections, but the throughput of these methods remains low despite recent advances in methodology (up to ∼50 neurons per single cortical area) (Economo et al., 2016; Gong et al., 2016; Guo et al., 2017; Lin et al., 2018; Zheng et al., 2013). Furthermore, these methods usually rely on dedicated imaging platforms specifically designed for neuronal tracing, which are not widely accessible to many laboratories.

To allow high-throughput projection mapping, we previously developed MAPseq (Han et al., 2018; Kebschull et al., 2016a) (Fig. 1A, *left*), a sequencing-based method capable of mapping projections of thousands of single neurons in a single brain. MAPseq achieves multiplexing by uniquely labeling individual neurons with random RNA sequences, or “barcodes.” However, like most other sequencing methods, the original MAPseq protocol relies on tissue homogenization, so the precise location of each soma is lost. Thus with MAPseq, it is difficult to combine information about projections with other types of information, such as gene expression and neuronal activity.

**Fig. 1.**
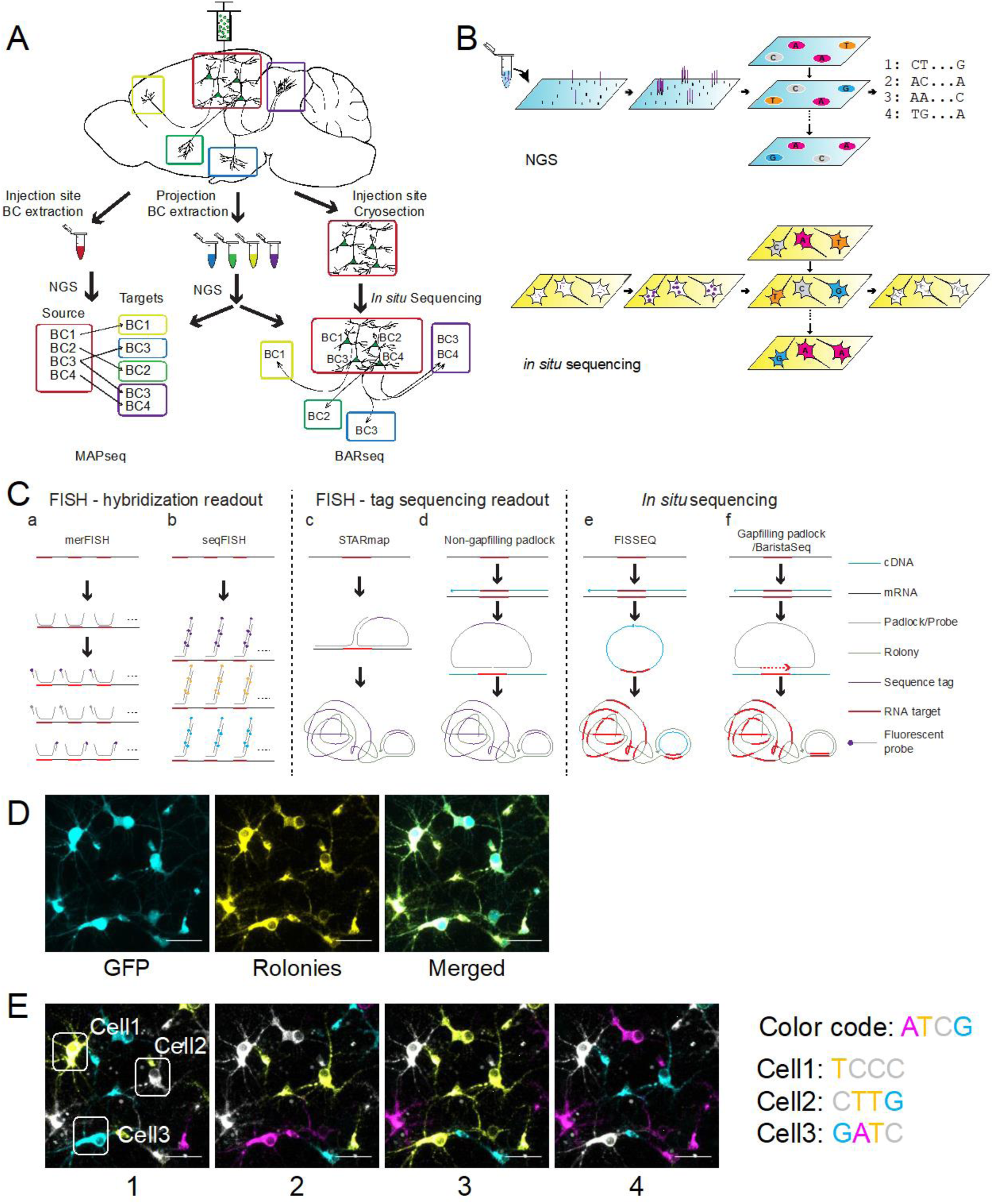
Multiplexed projection mapping using *in situ* sequencing. (A) Workflow of MAPseq (*left*) and BARseq (*right*). In both MAPseq and BARseq, a barcoded viral library is delivered to the area of interest. The source area and several target areas are then dissected. In MAPseq, barcodes in all dissected areas are sequenced using Next-Gen Sequencing (NGS). Barcodes at the source site are then matched to those at the targets to find the projection patterns of individual neurons. In BARseq, the injection site is sequenced *in situ*, thus preserving spatial information. (B) NGS vs. *in situ* sequencing. In conventional NGS (*top*), DNAs are anchored to a flow cell (blue), amplified locally, and sequenced. During each sequencing cycle, one fluorescence-labeled base is incorporated using the amplified DNA as template. The sequence is read out from the sequence of fluorescence over cycles. In *in situ* sequencing (*bottom*), RNA is reverse transcribed and amplified *in situ* on a slide (yellow). The amplified cDNAs are then sequenced using the same chemistry as NGS and imaged *in situ*. Sequences can be read out together with spatial information. (C) Comparison of *in situ* sequencing and hybridization techniques. Methods shown on the left use multiple rounds of hybridization to probe and read out multiple mRNAs. Methods shown in the middle use sequencing to multiplex read out of *in situ* hybridization signals. Methods shown on the right copy target sequences from the mRNA into the rolonies, allowing true sequencing of mRNA *in situ*. (a) In merFISH, multiple probes with barcode sequences on both ends are hybridized to each target mRNA. The probes are then read out using multiple rounds of hybridization using fluorescently labeled secondary probes. (b) In seqFISH, multiple rounds of HCR (hybridization chain reaction)-amplified FISH are carried out on the same target mRNAs, and the mRNA identities are read out using the color sequence. (c) In STARmap, a linear probe and a padlock probe are hybridized to adjacent sequences on the target. When bound, the linear probe acts as a splint to allow circularization of the padlock probe. The circularized padlock probe is then used as template for rolling circle amplification (RCA). (d) In the non-gapfilling padlock method, the target mRNA is reverse-transcribed to cDNA. A padlock probe then hybridizes to the cDNA and is circularized. The circularized padlock probe is then used as template for RCA. (e) In the Circligase-based FISSEQ, the cDNA is circularized using an ssDNA ligase. This circularized cDNA is then used as template for RCA. (f) The gapfilling padlock method is similar to the non-gapfilling version, except that a gap between the two arms of the padlock is filled by copying from the cDNA, thus allowing the actual cDNA to be sequenced. (D) Representative images of barcode rolonies (yellow) generated in primary hippocampal neuronal culture coexpressing barcodes and GFP (cyan). All GFP positive neurons were filled with barcode amplicons, indicating efficient barcode amplification in neuronal somata. (E) Images of the first four sequencing cycles of the same neurons shown in (D). The bases corresponding to the four colors and the sequences of the three neurons circled in (E) are indicated to the right. In all images, scale bars = 50 µm.

Here we describe BARseq (<u>B</u>arcoded <u>A</u>natomy <u>R</u>esolved by <u>Seq</u>uencing; Fig. 1A, *right*), a method that combines MAPseq with *in situ* sequencing (Chen et al., 2018; Ke et al., 2013; Lee et al., 2014) of cellular barcodes. BARseq preserves the spatial organization of neurons during projection mapping, and further allows co-registration of neuronal projections with mRNA expression in a single specimen. We show that BARseq is able to recapitulate the known spatial organization of major classes of neocortical excitatory neurons—corticothalamic (CT), pyramidal tract-like (PT-l; we call these neurons pyramidal tract-like because, unlike pyramidal tract neurons in the motor cortex, these neurons in the auditory cortex do not project to the spinal cord), and intratelencephalic (IT) neurons (Harris and Shepherd, 2015; Shepherd, 2013)—in the mouse auditory cortex and can additionally uncover previously unknown organization beyond these major classes. BARseq thereby complements traditional neuro-anatomical approaches by providing a high-throughput method for linking axonal projections, spatial organization, gene expression, and potentially other neuronal properties that define neuronal cell types and circuits.

## Results

In what follows, we first demonstrate robust *in situ* sequencing of cellular barcodes in neuronal culture. We then combine *in situ* barcode sequencing in cortical slices with mapping of projections (MAPseq). We next combine BARseq with *in situ* hybridization and subpopulation-targeted Cre labeling to correlate neuronal projections with gene expression and fluorescent labeling. Finally, we apply BARseq to understand the organization of projection neurons in the auditory cortex and identify a structured organization of projections beyond previously defined classes of projection neurons.

### *In situ* barcode sequencing in neuronal culture

We set out to develop a method capable of reading out highly diverse ensembles of sequences *in situ*, with cellular and even subcellular spatial resolution. Although several highly sensitive multiplexed *in situ* hybridization methods have been described [e.g., merFISH (Chen et al., 2015), seqFISH (Eng et al., 2019; Shah et al., 2016), STARmap (Wang et al., 2018), *in situ* sequencing (Ke et al., 2013) and osmFISH (Codeluppi et al., 2018)], these methods lack single-nucleotide specificity, and can distinguish only up to 10^2^-10^4^ different transcript sequences, each with a length of 10^2^-10^3^ nucleotides. By contrast, cellular barcodes used for projection mapping are much shorter (30-nt), more diverse (∼10^6^ to 10^7^), and may differ from each other by just a few nucleotides. Cellular barcodes thus cannot be distinguished using currently available multiplexed *in situ* hybridization approaches. Spatial transcriptomics (Rodriques et al., 2019; Stahl et al., 2016) can read out diverse short barcodes, but at present the spatial resolution is insufficient to resolve single neurons. Laser micro-dissection combined with RNA sequencing can provide cellular resolution, but the throughput is too low for high-throughput projection mapping. We therefore focused on an *in situ* sequencing approach (Chen et al., 2018; Ke et al., 2013) to achieve both the spatial resolution and the throughput needed for barcode identification *in situ*.

*In situ* sequencing is conceptually similar to conventional *in vitro* Illumina DNA sequencing (Fig. 1B), both of which consist of two basic steps: amplification and sequencing. In the amplification step, the RNA is converted to cDNA by reverse transcription, and the cDNA is amplified using rolling circle amplification, resulting in the formation of a small (< 1 µm) nanoball of DNA called a “rolony.” In the sequencing step, the rolonies are sequenced in parallel using four fluorescently-labeled nucleotides. The nucleotide sequences are thus transformed into color sequences, and are read out using multi-channel fluorescence microscopy.

To capture short diverse barcode sequences, we needed an approach in which the actual target barcode sequence is captured and amplified (Fig. 1Ca-f). Although some *in situ* hybridization methods use sequencing to read out gene-specific tags to allow multiplexed detection of RNAs (Ke et al., 2013; Wang et al., 2018), they cannot directly sequence the RNAs of interest and are unsuitable for barcode sequencing (Fig. 1Ccd). Circligase-based FISSEQ (Fig. 1Ce) can readout mRNA sequences, but is not sufficiently sensitive for barcode sequencing. We therefore used a targeted approach, BaristaSeq (Chen et al., 2018), in which reverse transcription was used to convert the barcode sequence into cDNA, after which padlock probe hybridization followed by gap-filling and ligation was used to form a circular template for rolling circle amplification (Fig. 1Cf). The gap-filling padlock approach allowed for efficient amplification of barcodes in neurons co-expressing barcodes and GFP (Fig. 1D). We also adapted Illumina Sequencing By Synthesis (SBS) chemistry for *in situ* reactions. In combination, these strategies formed the basis for a protocol that allowed efficient and robust sequencing of barcoded cultured neurons (Fig. 1E).

### *In situ* barcode sequencing in brain slices

*In situ* sequencing in brain slices presents additional challenges compared to sequencing in cultured neurons. We developed a chamber system which enabled convenient handling and was compatible with the amplification reactions (Fig. S1A; see STAR Methods). Because barcoded neurons express high levels of GFP (which interferes with imaging during sequencing), we developed a protocol for tissue digestion to reduce the GFP signal from barcoded cells (Fig. S1B; see STAR Methods) and to increase RNA accessibility. Together these optimizations resulted in highly efficient amplification of barcodes (Fig. S1C) compared to other methods (Ke et al., 2013; Lee et al., 2014). We also optimized Illumina sequencing chemistry for brain slices, resulting in a ∼10-fold improvement in signal-to-noise ratio (SNR) compared to sequencing by ligation approaches (Ke et al., 2013; Lee et al., 2014) (Fig. S1D, E; see STAR Methods). These modifications resulted in highly efficient and specific barcode amplification (Fig. 2A) and sequencing (Fig. 2B) in barcoded neuronal somata in brain slices.

**Fig. 2.**
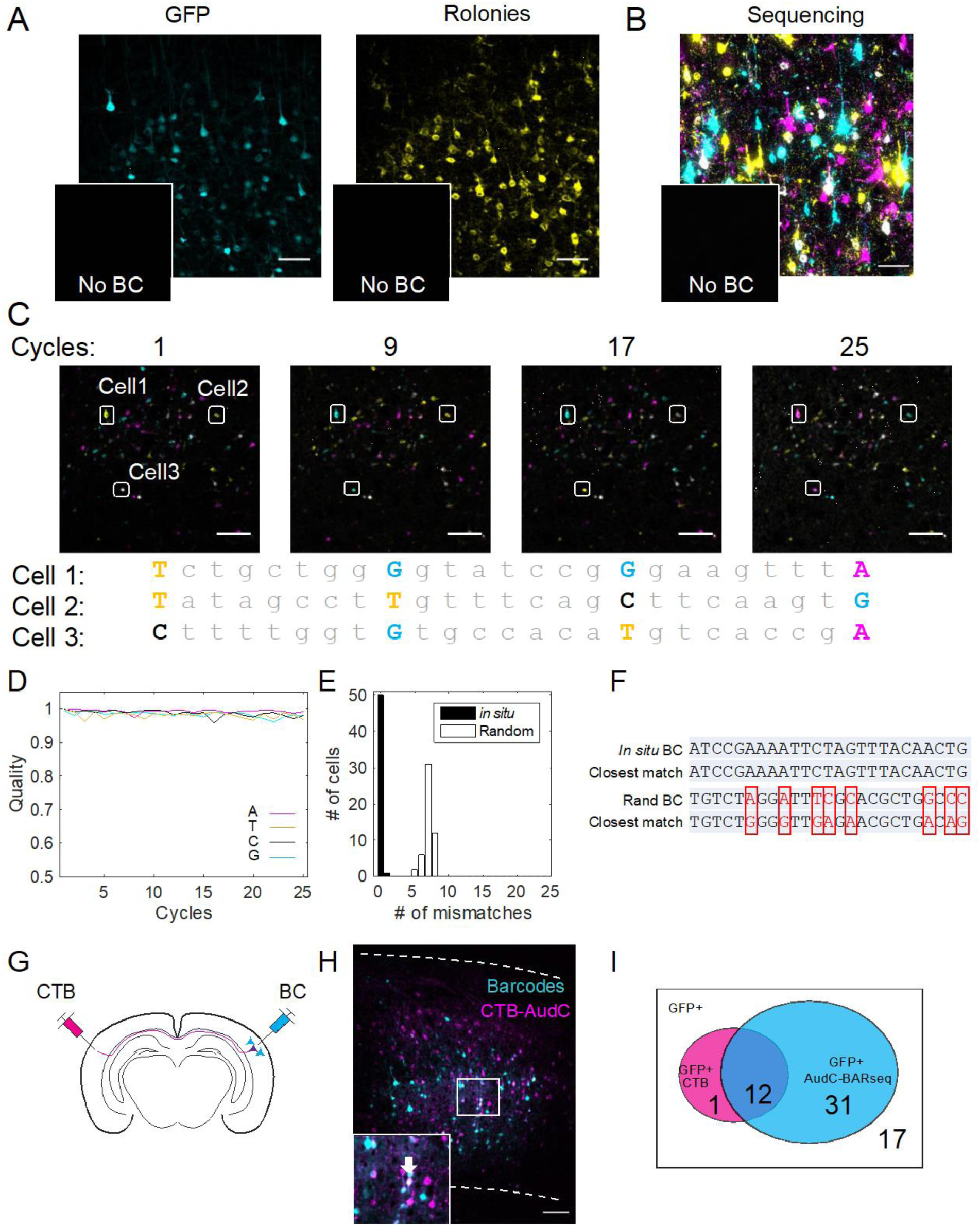
Validation of BARseq. (A) Representative low-magnification images of a barcoded brain slice expressing GFP (*left*) and rolonies (*right*) generated in the same brain slice. GFP intensity does not correlate perfectly with rolony intensity due to differences in protein and RNA expression. Insets: Negative control images of GFP and rolonies of a non-barcoded brain slice taken with the same exposure settings. No GFP or rolonies are visible in these images. Scale bars = 50 µm. (B) Representative high-resolution sequencing image of a barcoded brain slice. The color code is shown in Fig. 1E. Inset shows sequencing image of a non-barcoded brain slice. Scale bar = 50 µm. (C) Representative low-resolution images of the indicated cycles of barcode sequencing in a brain slice. The sequences of the three cells base-called are indicated below the images. Only bases corresponding to the images shown are capitalized and color-coded. Scale bars = 100 µm. (D) The quality of the base calls over 25 cycles of Illumina sequencing *in situ* on the barcoded brain slice. The quality score is defined as the intensity of the called channel divided by the root sum square of all four channels. A quality score of 1 (best) indicates sequencing signal in only one channel, and a score of 0.5 (worst) indicate same intensity across all four channels. (E) Histogram of the number of mismatches between the *in situ* reads and their closest matches from *in vitro* reads (*in situ*) and the number of mismatches between random sequences and their closest matches from *in vitro* reads (Random). (F) An example barcode read *in situ* and its closest match *in vitro*, and a random sequence and its closest match *in vitro*. Red indicates mismatches. (G) A brain was injected with CTB in the contralateral auditory cortex and barcoded in the ipsilateral auditory cortex. BARseq results of the barcoded neurons were then compared to retrograde labeling by CTB. (H) A representative image of a brain slice double labeled with barcodes (cyan) and CTB (magenta) from the contralateral auditory cortex. Dashed lines indicate the top and the bottom of the cortex. Scale bar = 100 µm. A magnified view of the squared area is shown in the inset. The arrow in the inset indicates a GFP+ CTB+ double-labeled neuron. (I) Venn diagram showing the number of GFP expressing neurons labeled with (magenta) or without (white) CTB and/or neurons found to project contralaterally using BARseq (cyan). See also Fig. S1 and Table S1.

To evaluate barcode sequencing in brain slices, we sequenced 25 bases in a sample infected with a diverse (>10^6^ unique sequences) pool of barcoded Sindbis virus (Kebschull et al., 2016a; Kebschull et al., 2016b) injected in the auditory cortex (Fig. 2C). Basecall quality (Fig. 2D; Fig. S1F; see STAR Methods) and signal intensity (Fig. S1G) remained high through all 25 cycles. We observed no bias toward any particular base (Fig. S1H). Read accuracy was high: Barcodes in 50/51 (98%) randomly selected cells matched perfectly to known barcodes in the library, and the remaining cell had only a single base mismatch to the closest barcode in the library (Fig. 2E, F). The per-base error rate of 1 / (25 × 51) = 0.0078 corresponds to a Phred score of 31. Because the barcodes in this library represent a small fraction (∼10^−9^) of all possible 25-mer barcodes (>10^15^), random 25-nt sequences on average had seven mismatches to the closest known barcode, indicating that our barcode reads were unlikely to be false positive matches by chance. These results indicate that *in situ* sequencing of RNA barcodes in brain slices is both accurate and efficient.

### Projection mapping using BARseq

We next combined *in situ* barcode sequencing with MAPseq. In MAPseq (Fig. 1A, *left*), neurons are barcoded using a Sindbis virus library. Both the source area containing neuronal somata and target projection areas are micro-dissected into “cubelets” and sequenced. Barcodes from the target areas are then matched to those at the source area to reveal projection patterns. The spatial resolution of MAPseq is thus determined by the size of the cubelets. In BARseq, we perform *in situ* sequencing of barcoded somata at the source; the target projection areas are still micro-dissected and sequenced as cubelets (Fig. 1A, *right*). BARseq thus inherits the throughput and cubelet resolution of projections of MAPseq, but allows the precise somatic origin of each axonal projection to be determined with cellular resolution.

The high sensitivity and accuracy of MAPseq is well established (Han et al., 2018; Kebschull et al., 2016a; Klingler et al., 2018). In particular, the sensitivity of MAPseq is indistinguishable from that of conventional GFP-based single neuron reconstruction (Han et al., 2018). To confirm that BARseq maintains the high sensitivity and accuracy of MAPseq, we compared it to conventional retrograde cholera toxin subunit B (CTB) tracing of contralateral projections. We injected a highly diverse (∼10^7^ barcodes) viral library into the auditory cortex and CTB in the contralateral auditory cortex (Fig. 2G, H). We used BARseq to identify contralaterally projecting neurons and the precise locations of their somata. We then identified barcoded somata that were also labeled with CTB. The majority (12/13, 92%) of CTB-labeled neurons were detected by BARseq, whereas only 28% (12/43) of contralaterally projecting neurons identified by BARseq were labeled with CTB (Fig. 2I; Table S1). The strength of the contralateral projections was well above the noise threshold (defined by barcode counts in the olfactory bulb, an area to which the auditory cortex does not project; Fig. S1I), indicating that the higher apparent sensitivity of BARseq was not due to false positives resulting from contaminating barcodes. These results indicate that BARseq retains the previously reported (Kebschull et al., 2016a) high specificity and sensitivity of MAPseq (91 ± 6%), and may exceed the sensitivity of conventional CTB tracing.

### Relating projection patterns to gene expression using BARseq

The *in situ* nature of BARseq allows us to relate axonal projection patterns with other single neuron characteristics measured *in situ* in the same sample. In particular, BARseq can relate projections to past or current gene expression by combining with marker-based *Cre* driver lines or *in situ* hybridization.

To demonstrate BARseq in Cre-labeled neuronal subpopulations, we performed BARseq in the auditory cortex in a Fezf2-2A-CreER::Rosa-CAG-LSL-tdTomato (Ai14) mouse induced at age 6 and 7 postnatal weeks (Fig. 3A). Fezf2 is predominantly expressed in layer 5 PT-l, but is also expressed at lower levels in some layer 5 IT neurons and layer 6 CT neurons (Tasic et al., 2018). Projection patterns obtained from bulk tracing were mostly those expected from PT-l neurons, but also included some callosal projections, suggesting a small contribution from Fezf2-expressing IT neurons (Matho et al., in preparation). We sequenced 1291 projection neurons with good sequencing quality *in situ,* including 72 neurons co-labeled by tdTomato driven by Fezf2-2A-CreER, and assayed their projection patterns to 11 target areas (see STAR Methods). The Fezf2+ population included 53 (74%) PT-l neurons projecting to the tectum and the thalamus, eight (11%) CT neurons projecting to the thalamus only, nine (13%) IT neurons in layer 5 projecting to the contralateral cortex, and two (3%) neurons in layer 5 projecting to the striatum (Fig. 3B, C). The lack of corticotectal projection in the callosal projecting neurons is unlikely an artifact caused by limited detection sensitivity of corticotectal projections, because these neurons also project to other telencephalic areas that PT-l neurons do not. Neurons labeled with Fezf2-2A-CreER thus consisted mainly of PT-l neurons, along with a small fraction of callosally-projecting neurons in layer 5 and CT neurons in layer 6. The projection patterns of Fezf2+ neurons were thus consistent with previous bulk cell-type specific tracing results using the same Cre line (Matho et al., in preparation). These results demonstrate the combination of BARseq with Cre-dependent labeling of neuronal subpopulations, and further validate the results of BARseq.

**Fig. 3.**
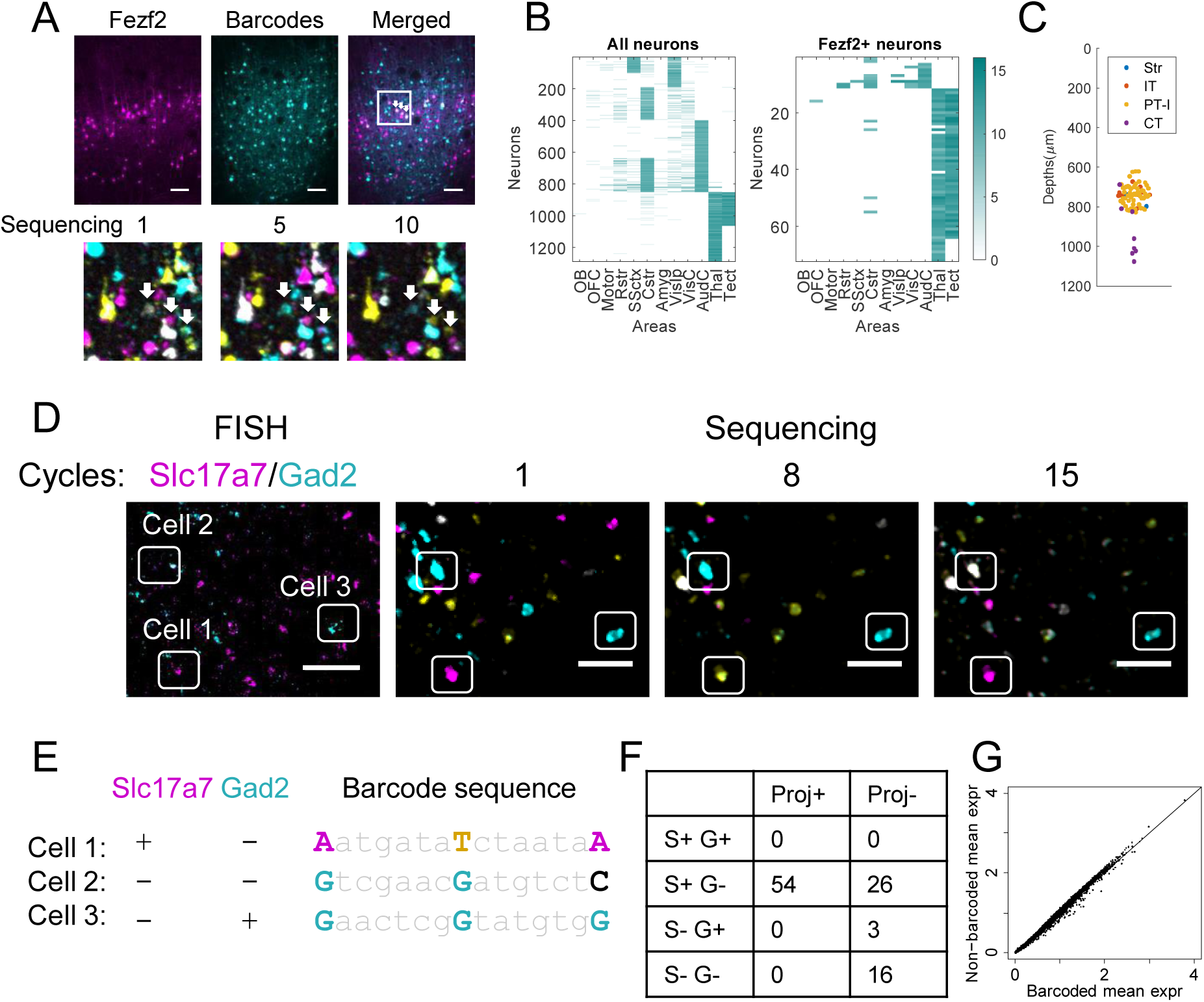
Correlating gene expression and projections using BARseq. (A) Barcode sequencing in Fezf2-2A-CreER::Rosa-CAG-loxp-STOP-loxp-tdTomato (Ai14) animals. Top row: images of Fezf2+ tdTomato expressing cells driven by Fezf2-2A-CreER::Ai14 (left), GFP expressing barcoded cells (middle), and merged image of the two (right). Scale bar = 100 µm. Bottom row: sequencing images of the indicated cycle of the area indicated in the merged image. The three arrows indicate tdTomato-expressing barcoded neurons. (B) Projection patterns of all neurons (left) and Fezf2+ neurons (right). Rows indicate neurons and columns indicate projection areas. (C) The cortical depth distribution of barcoded Fezf2-2A-CreER::Ai14 labeled somata. Neurons are color-coded by projections as CT (thalamic projections), PT-l (tectal and thalamic projections), IT (callosal projections), and Str (striatal projections only). (D)(E) Representative high-resolution image of FISH (left) against Slc17a7 (magenta) and Gad2 (cyan), and low-resolution sequencing images of cycle 1, 8, and 15 of the same sample are shown in (D). The three cells circled in (D) are base-called in (E) with their expression profile of Slc17a7 and Gad2 shown. Scale bars = 50 µm. (F) The number of neurons with (Proj+) or without (Proj-) projections in excitatory neurons (S+ G-), inhibitory neurons (S- G+), and non-neuronal cells (S- G-). No cell was found expressing both Slc17a7 and Gad2 (S+ G+). (G) Mean log normalized expression of each gene averaged over all barcoded (x-axis) and non-barcoded (y-axis) neurons. The gene expression is regressed with a Poisson model to remove the effect of both the percentage of mitochondrial genes and endogenous UMI counts. The diagonal line indicates equal expression in barcoded vs. non-barcoded cells. See also Fig. S2 and Table S2.

To correlate projections with the expression of multiple genes, BARseq can be combined with *in situ* detection of genes in the same sample. To demonstrate the feasibility of resolving both barcodes (for projection mapping) and endogenous gene expression *in situ*, we combined BARseq with fluorescent *in situ* hybridization (FISH) against Slc17a7 [a marker of excitatory neurons; (Tasic et al., 2016)] and Gad2 (a marker of inhibitory neurons) in the same cells (Fig. 3D, E). Consistent with the fact that most projection neurons in the cortex are excitatory, we identified 54 neurons with long-range projections, all of which expressed Slc17a7, but not Gad2 (Fig. 3F; Table S2; see STAR Methods for details). The projection neurons identified by BARseq were thus consistent with their transcriptional cell types. These experiments demonstrate that projection mapping with BARseq is compatible with *in situ* interrogation of gene expression.

We next assessed whether barcoding disrupted the transcriptomic landscape of single neurons. We performed single-cell RNAseq in 398 barcoded and 1088 non-barcoded cells dissociated from the mouse auditory cortex. Although fewer endogenous mRNA molecules were recovered from barcoded compared to non-barcoded neurons, the relative levels of endogenous gene expression remained largely unchanged in barcoded cells (Fig. 3G; Fig. S2A; see STAR Methods for details). The fact that relative levels were preserved is compatible with a model in which infection with the barcoded viral vector affects the expression of most endogenous genes about equally. Importantly, top principal components of genes identified in the non-barcoded neurons were equally effective in describing gene expression in barcoded neurons (Fig. S2B). These principal components contained many known neuronal subtype markers (Fig. S2C). These results indicate that the expression pattern of genes, especially those that contribute to cell typing, remain largely intact in barcoded cells.

### Projection diversity in the mouse auditory cortex

Having established the sensitivity and specificity of BARseq, we applied it to study the organization of long-range projections from the mouse auditory cortex. We performed BARseq on two brains (XC9 and XC28) and MAPseq on an additional brain (XC14). We allowed one mismatch when matching *in situ* barcodes to the projection barcodes (see STAR Methods; Fig. 4A; Fig. S3A), and found 1806 (55%) of 3237 cells sequenced *in situ* projected to at least one target area (Table S3). We further excluded cells obtained from tissue deformed during processing, and from labeled cells outside of the cortex. This resulted in 1309 neurons sequenced *in situ* and 5082 neurons sequenced *in vitro* using conventional MAPseq (Table S3; Fig. 4B). Because only neurons at the center of the injection sites were sequenced in each BARseq brain, the MAPseq experiment produced more neurons per brain than BARseq, but might include neurons from nearby cortical areas.

**Fig. 4.**
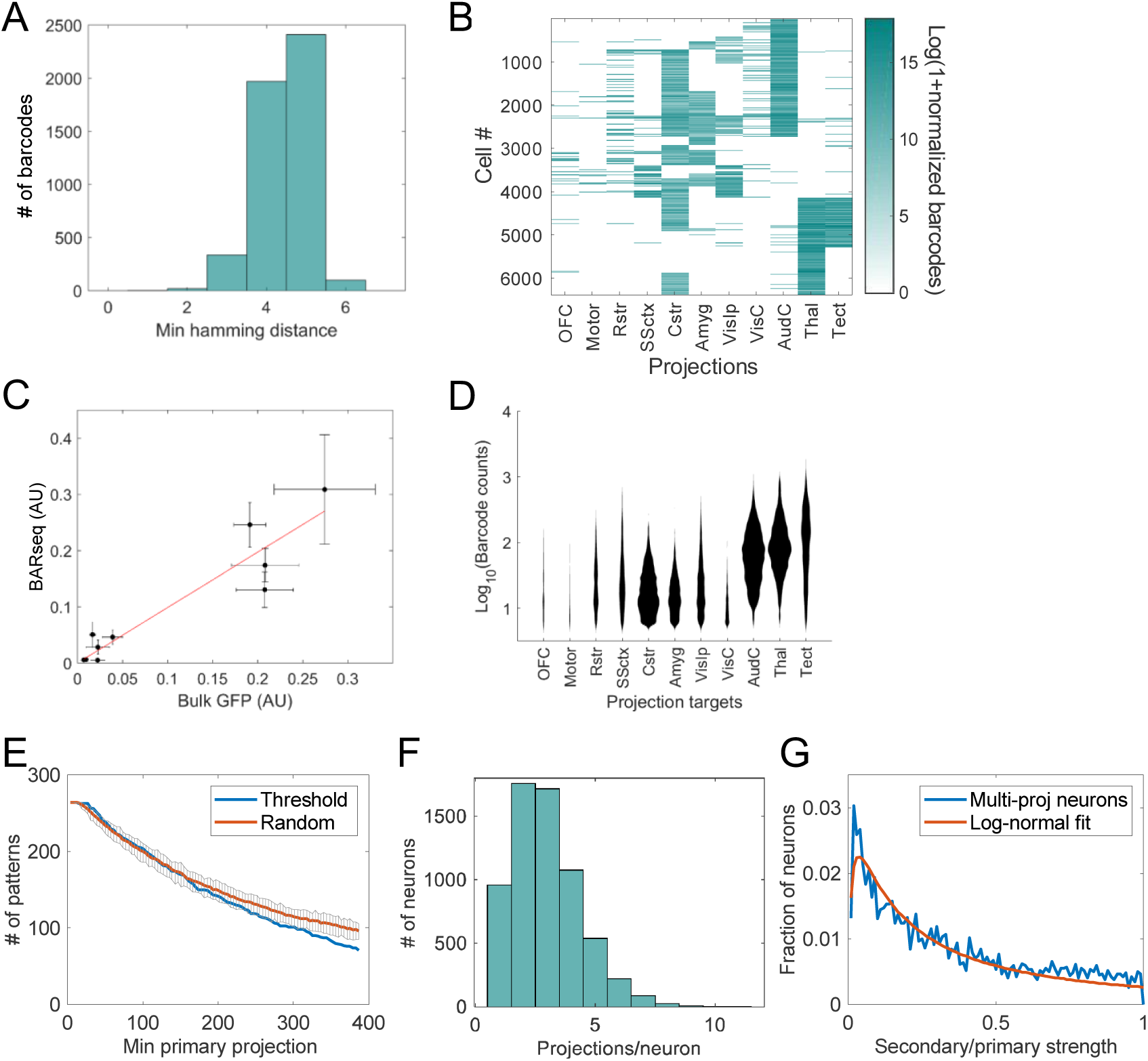
Mapping projections of the auditory cortex using BARseq. (A) Histogram of the minimal pairwise hamming distance of the first 15 bases of barcodes recovered from brain XC9. (B) Single-cell projection patterns sorted by clusters. Each row represents a barcode and each column represents projection strengths to the indicated brain area. OFC: orbitofrontal cortex; Motor: motor cortex; Rstr: rostral striatum; SSctx: somatosensory cortex; Cstr: caudal striatum; Amyg: amygdala; VisIp: ipsilateral visual cortex; VisC: contralateral visual cortex; AudC: contralateral auditory cortex; Thal: thalamus; Tect: tectum. (C) Conventional bulk GFP tracing intensities (x-axis) were plotted against the bulk projection strength obtained from BARseq (y-axis). Error bars indicate SEM. N = 5 for GFP tracing and N = 3 for BARseq. Pearson correlation coefficient r = 0.94, p < 0.0001. (D) The distribution of projection intensity in each projection area. The y-axis indicates the logarithms of raw barcode counts in each area, and the x-axis indicates the number of cells. (E) The numbers of binarized projection patterns (y-axis) after filtering for primary projection strength (x-axis). Blue line indicates filtering at the indicated thresholds, and red line indicates random subsampling to the same sample size. Black lines and error bars indicate 95% confidence interval for subsampling. (F) Histogram of the number of projections per neuron. (G) The fractions of multi-projection neurons (y-axis) are plotted against the ratio between the secondary and primary projections (x-axis). Blue line indicates actual distribution and red line indicates fitting with a log normal distribution. See also Fig. S3 and Table S3.

We focused on 11 auditory cortex projection target areas, including four ipsilateral cortical areas (orbitofrontal, motor, somatosensory, and visual), two contralateral cortical areas (visual and auditory), three subcortical telencephalic areas (amygdala, rostral striatum, and caudal striatum), the thalamus, and the tectum (see STAR Methods for details of dissected areas). These included most major brain areas to which the auditory cortex projects, as determined by conventional bulk GFP tracing experiments (Oh et al., 2014). We also collected tissue from the olfactory bulb, an area to which the auditory cortex does not project, as a negative control.

Only five out of 6391 neurons had non-zero barcode counts in the olfactory bulb (four neurons with a count of one molecule, and one neuron with a count of four molecules), indicating that the technical false-positive rate for a single target area is extremely low at 5 / 6391 = 0.08%. Using the false-positive rate and the number of barcodes with zero counts in each area, we estimated that a total of 40 out of 18851 projections detected in our dataset might be false positives, corresponding to a false-discovery rate of 0.2% (i.e., for each 1000 detected projections, two were likely false positives). BARseq thus has a very low false discovery rate.

MAPseq has now been validated using several different methods, including single neuron reconstruction, in multiple brain circuits (Han et al., 2018; Kebschull et al., 2016a). The contribution of potential artifacts, including those due to non-uniform barcode transport and variable barcode expression strength, have been carefully quantified in previous work (Han et al., 2018; Kebschull et al., 2016a). Although one potential challenge of MAPseq arises from fibers of passage, in practice these have not represented a major source of artifact (Han et al., 2018; Kebschull et al., 2016a). One reason is that the total amount of axonal material due to fibers of passage is typically small compared with the rich innervation of a fiber at its target. A second reason is that the barcode carrier protein is designed to be enriched at synapses (Kebschull et al., 2016a), which further minimizes the contribution of passing fibers to barcode counts. To determine the extent of contamination from fibers of passage, we examined the striatal projections of putative CT neurons (i.e. neurons that project to the thalamus, but not to any cortical area or the tectum) and putative PT-l neurons (i.e. neurons that project to the tectum, the thalamus, but not to any cortical area) from the two BARseq brains (the MAPseq brain was excluded because it may include PT-l neurons from neighboring areas that did not project to the tectum, and may be misidentified as CT neurons). Both types of neurons send descending fibers through the internal capsule adjacent to the caudal striatum, and thus can be used to evaluate fibers of passage contamination in the striatum. Consistent with the fact that PT-l but not CT neurons project to the striatum, only 8 / 250 (3.2%) putative CT neurons sequenced *in situ* projected to the caudal striatum, whereas 59 / 245 (24%) putative PT-l neurons sequenced *in situ* projected to the caudal striatum. The fact that ∼97% of putative CT neurons did not show projection to the caudal striatum reinforce previous observations that fibers of passage have minimal impact on projections mapped in these experiments.

At the mesoscopic level, the projection patterns of the auditory cortex revealed by BARseq were consistent with anterograde tracing in the Allen Mouse Connectivity Atlas (Oh et al., 2014). In MAPseq and BARseq, the strength of the projection of a neuron to a target is given by the number of individual barcode molecules from that neuron recovered at the target. This is analogous to a conventional fluorophore mapping experiment, in which projection strength is assumed to be proportional to GFP intensity. Consistent with conventional bulk GFP tracing (Oh et al., 2014) (Fig. 4C; Pearson correlation coefficient r = 0.94, p < 0.0001), projections from the auditory cortex to the thalamus, the tectum, the contralateral auditory cortex, and the caudal striatum were particularly strong (Fig. 4D). BARseq thus provides quantitative measurements of long-range projections in the auditory cortex consistent with those obtained by conventional bulk labeling techniques.

The projection patterns of individual neurons were remarkably diverse. To quantify this diversity, we binarized projection patterns, using a conservative threshold for projection detection (four molecules per barcode per area, given by the OB negative control). We observed 264 distinct patterns, or 13% of the 2^11^ = 2048 possible patterns. This high diversity was unlikely an artifact of projections missed by BARseq due to false negatives: 237 to 247 unique projection patterns remain even when a false negative rate of 10-15% was assumed (see STAR Methods). This high diversity was also unlikely to be caused by limited sensitivity of MAPseq for neurons with fewer barcodes overall: A significant fraction of these projection patterns (112 / 264 = 42%) can be accounted for solely by strong projections (see STAR Methods), and eliminating neurons with weaker primary projections resulted in only a moderate reduction in the diversity of projection patterns (Fig. 4E, blue line). Furthermore, this reduction was largely due to the reduction in sample size (Fig. 4E, red line). Because of the conservative thresholding we used and the fact that the auditory cortex may project to areas we did not sample, the actual number of distinct projection patterns may be higher.

The majority (85%) of cortical projection neurons projected to two or more target areas (Fig. 4F) after binarization. The ratio of the projection strengths of the two strongest projections follow a log normal distribution (Fig. 4G), and about half of all neurons have a secondary projection that is at least 20% as strong as their primary projections. These estimates were based on neurons whose primary projections were strong enough to observe a secondary projection, and therefore were not limited by the sensitivity of MAPseq/BARseq or by the variability in the expression level of barcodes. This high projection diversity and multiple projections per neuron are consistent with findings in the visual cortex (Han et al., 2018), and may be a general feature of cortical projections.

### BARseq recovers structured projections across major classes of projection neurons

Simple hierarchical clustering on the non-binarized projections (see STAR Methods) partitioned neurons into established (Harris and Shepherd, 2015) cortical classes: CT (corticothalamic), PT-l (pyramidal tract-like), and IT (intratelencephalic, Fig. 5A). CT neurons project to the thalamus; PT-l neurons project to both the thalamus and the tectum; and IT neurons project to the cortex but not subcortically to the thalamus or tectum. IT neurons were further subdivided into those that project to both the ipsilateral and contralateral cortex (the ITc class), and those that project ipsilaterally only (the ITi class). These patterns correspond to the classically-defined top-level subdivisions of cortical projection patterns.

**Fig. 5.**
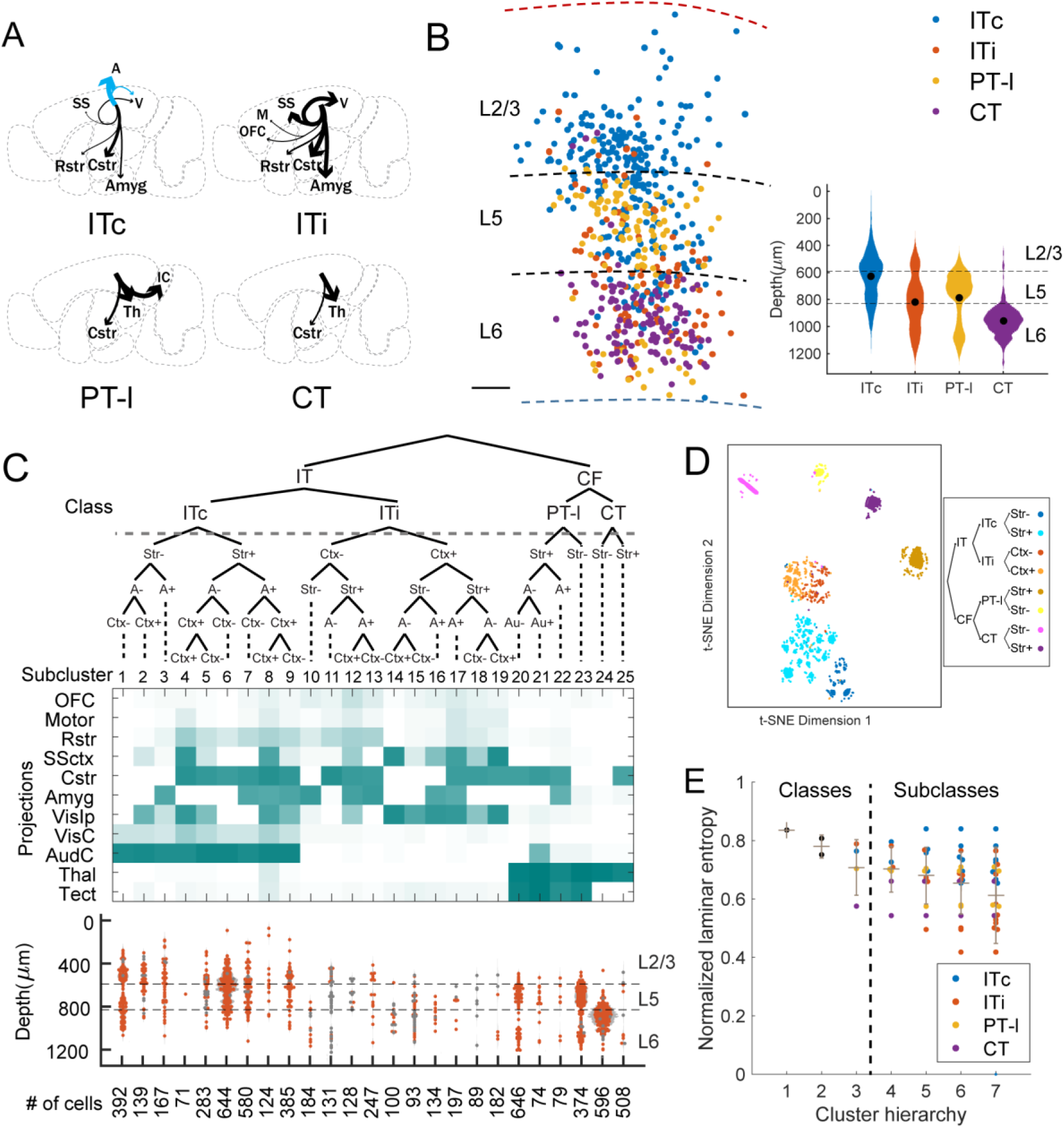
The laminar organization of projection neurons in the auditory cortex. (A) The mean projection patterns of clusters corresponding to the indicated major classes of neurons. Line thickness indicates projection strength normalized to the strongest projection for that class. Blue arrows indicate projections to contralateral brain areas and black arrows indicate projections to ipsilateral brain areas. (B) The sequenced projection neurons from a brain (XC9) are color-coded by class identities and plotted at their locations in the cortex. The top and bottom of the cortex are indicated by the red and blue dashed lines, respectively. The laminae and their boundaries are marked. Scale bar = 100 µm. Inset: histograms of each class of projection neurons in the pooled BARseq dataset. The y-axis indicates cortical depth and the x-axis indicates the number of neurons at that depth in a particular class. The laminae are indicated on the right and the boundaries between two layers are marked by dashed lines. (C) Hierarchical clustering of single-cell projection data. *Top*: dendrogram of the hierarchical structure of the clusters. An index is assigned to each leaf as indicated. Nodes representing major classes are labeled by class names, and nodes representing finer divisions are labeled by major differences in projections. *Middle*: the mean projection patterns of the corresponding leaf cluster. Each column represents a leaf cluster. Each row represents projection to the indicated area as labeled on the left. *Bottom*: The laminar distribution of the corresponding leaf clusters. Individual neurons are superimposed on top of the distribution plots (light grey). Neurons whose cluster identity were less confident were marked in gray. The number of cells that belong to each leaf cluster is indicated below. Neurons of subcluster 25 were likely misidentified PT-l neurons (see STAR Methods). (D) t-SNE plot of the projection neurons. The neurons are color-coded by their first level subcluster identities post-hoc. (E) The normalized entropy of nodes/leaves (y-axis) in the indicated clustering hierarchy (x-axis). The branch of the subcluster nodes/leaves were color coded as indicated. Grey bars indicate mean ± stdev of all nodes/leaves of a specific hierarchy. Hierarchies 1-3 correspond to class divisions and hierarchies 4-7 correspond to subcluster divisions. See also Fig. S4-S6.

The classification of cortical excitatory projection patterns into PT-l, IT and CT classes was supported by the corresponding segregation of these neurons into distinct laminae. As expected, the major classes segregated strongly with laminar position: Superficial layers contained predominantly IT neurons, whereas deep layers contained predominantly PT-l and CT neurons (Harris and Shepherd, 2015) (Fig. 5B; see STAR Methods and Fig. S3B-E for the estimation of layers using layer-specific marker genes). Thus major classes of cortical projection neurons defined by BARseq had distinct laminar distributions, consistent with previous observations using conventional methods (Harris and Shepherd, 2015).

Our data revealed two interesting features of the PT-l neurons in the auditory cortex. First, although IT neurons projected to both the caudal striatum—an area implicated in auditory decision making (Znamenskiy and Zador, 2013)—and the rostral striatum, PT-l neurons projected only to the caudal striatum. We confirmed this finding by triple retrograde tracing (Fig. S4A): A significant fraction of neurons (66 / 296) projecting to the rostral striatum also projected to the caudal striatum, but none (0 / 296) projected to the tectum (Fig. S4B, C). Second, we identified a minor population of PT-l neurons in layer 6 of the auditory cortex in addition to the more common layer 5 PT-l neurons (Fig. 5B). This population of layer 6 PT-l neurons has previously been reported to differ from the layer 5 PT-l neurons based on physiological and other properties (Slater et al., 2013). Similar to the differences observed between the layer 5 and layer 6 PT-l neurons, the corticotectal projections were slightly stronger in layer 5 than those in layer 6, but with considerable overlap [Fig. S4D (PT-Tect); 11.7 ± 2.2 (mean ± stdev), N = 80 for layer 6 neurons and 13.8 ± 1.6, N = 185 for layer 5 neurons, p < 0.0001 using bootstrap K-S test]. The strengths of other major projections were indistinguishable between the two groups (Fig. S4D-PT, p = 0.5 for both corticothalamic projections and corticostriatal projections using bootstrap K-S test). The results of BARseq were thus consistent with, but went beyond, classical taxonomy.

### Projection subclusters do not segregate by laminae

Even within the major classes, projections from the auditory cortex to individual targets were often correlated (Fig. S5A). For example, an ITi neuron with a strong projection to the somatosensory cortex was also likely to have a strong projection to the visual cortex, but a weak striatal projection (Fig S5Ab). Such correlations suggest that the remarkable diversity of projections did not arise purely by chance, e.g. by a process in which each neuron selected targets at random. We therefore exploited the large sample size available from BARseq to look for statistical signatures of the organization of projections from auditory cortex.

To explore the statistical structure underlying these projection patterns, we performed hierarchical clustering on the projection patterns. For this analysis, we did not binarize the projection patterns; we considered not only whether a neuron projected to an area, but also the strength of that projection. This clustering uncovered structure well beyond the previously established top-level classes (Fig. 5C, D; see Fig. S5B-E and STAR Methods for details of clustering methods). Depending on the precise clustering algorithm and statistical criteria used (Fig. S5F), as many as 25 subclusters were revealed. Most of these subclusters were robust: 5682/6391 (89%) of neurons were unambiguously assigned into one of the 25 subclusters using a probabilistic approach (see STAR Methods; Fig. S5G, H). All top-level subclusters were observed in all three brains (Fig. S6A; see STAR Methods for likely misclassification of some striatal-projecting PT-l neurons as CT neurons due to choice of projection targets sampled). BARseq thus uncovered organization in projections well beyond the previously described major divisions.

As noted above, the three major cortical classes—PT-l, IT and CT—are spatially segregated into laminae. By contrast, most subclusters defined by projection pattern, especially those within the IT class, spanned many layers (Fig. 5C, E; p = 0.95 for cluster hierarchy 2-7, Kurskal-Wallis test; see STAR Methods). For example, a subcluster consisting of IT neurons that project only to contralateral cortical areas, but not the ipsilateral cortex or the striatum (Leaf 1, Fig. 5C), can be found across all laminae. This mixed laminar distribution of projection patterns was consistent across brains (Fig. S6B, C; see STAR Methods). Therefore, laminar position could not fully explain projection patterns or diversity.

### BARseq associates projections with transcriptionally defined IT subtypes

Although projection subclusters did not segregate cleanly into laminae, we hypothesized that the conjunction of laminar position and gene expression might predict projection patterns. Although gene expression patterns of single neurons have been studied in other cortical areas, including visual, somatosensory, and motor cortices (Tasic et al., 2018; Zeisel et al., 2015), gene expression patterns of auditory cortical neurons have not previously been explored. We therefore performed single-cell RNAseq for 735 neurons in the auditory cortex using 10x Genomics Chromium. As expected from work in other cortical regions, the top-level partition among neurons was between excitatory and inhibitory classes (Fig. 6A). Within the excitatory neuron class, gene expression further segregated into CT and IT subtypes. As in other studies using the 10x platform (Zeisel et al., 2018), few PT-l neurons were recovered, possibly due to biases in cell survival (see STAR Methods). The transcriptional taxonomy of neurons in the auditory cortex were thus consistent with other sensory cortices.

**Fig. 6.**
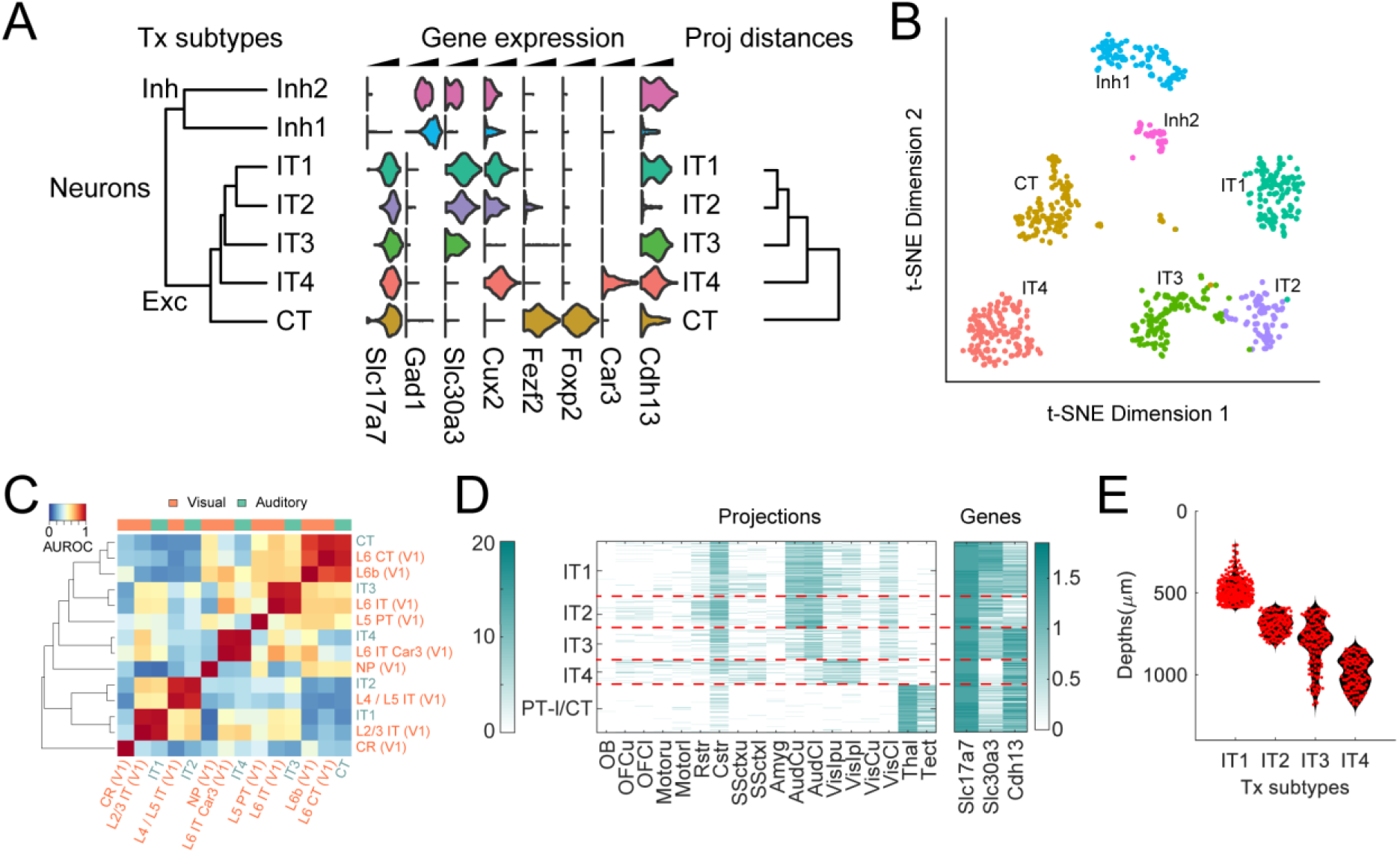
Subtypes of IT neurons defined by gene expression in the auditory cortex. (A) Histograms of the log normalized expression of the indicated marker genes in the indicated clusters obtained from single-cell RNAseq in the auditory cortex. The dendrogram on the left shows distances of mean gene expression among transcriptomic clusters, and the dendrogram on the right show distances of mean projection pattern among the same subtypes of neurons obtained through BARseq and FISH. (B) t-SNE plot of the gene expression of neurons color-coded by cluster identity as in (A). (C) MetaNeighbor comparison of neuronal clusters obtained in the auditory cortex to those in the visual cortex from Tasic et al. (2018). (D) Projections (left) and the expression of genes (right) of neurons obtained using combination of BARseq and FISH are shown on a log scale. The transcriptionally defined subtype labels are shown on the left. The color bar on the left indicates Log2 values of normalized barcode counts and the color bar on the right indicates Log10 values of normalized mRNA counts. Projection areas are the same as in Fig. 4B, except that each cortical area is divided into upper (u) and lower (l) layers. (E) Distributions of laminar positions of neurons. Individual neurons (red) are superimposed on the smoothed distribution (black). See also Fig. S7.

To further explore whether transcriptionally-defined subtypes corresponded to subclusters defined by projection pattern, we focused on subdivisions within the IT class. IT neurons were the most diverse class based on projection mapping, with as many as 19 distinct subclusters (Fig. 5C). Clustering of gene expression revealed four top-level IT subtypes, which we denote IT1, IT2, IT3, and IT4 (Fig. 6A, B). These four subtypes corresponded well to the four major subtypes of IT neurons previously identified in visual cortex (Tasic et al., 2018)(Fig. 6C; see STAR Methods). We thus sought to determine how the 19 projection subclusters were partitioned among the 4 transcriptionally defined subtypes.

To assess the relationship between projection pattern and gene expression among IT neurons at the single neuron level, we combined BARseq with fluorescence *in situ* hybridization (FISH) against Cdh13, Slc30a3, and Slc17a7 (Fig. 6D). The combination of laminar position and expression of these markers allowed us to assign neurons to subtypes: IT1 neurons consisted of all IT neurons in layer 2/3; IT2 neurons consisted of Cdh13-negative IT neurons in layer 5; IT3 neurons consisted of both Cdh13-expressing neurons in layer 5 and Slc30a3-expressing neurons in L6; and IT4 neurons consist of Slc30a3-negative IT neurons in layer 6 cortex (Tasic et al., 2018)(Fig. 6D, E; see STAR Methods). We collected the same projection areas as in the previous BARseq experiments, but achieved higher spatial resolution of the projection targets by collecting upper and lower cortical layers separately. We obtained good sequencing and FISH in 979 projection neurons from two brains (Fig. 6D; Fig. S7A), of which 735 were IT neurons defined by projection (note that this set of 735 IT neurons was independent of the 735 neurons studied using single-cell RNAseq; the same number of neurons is coincidental). These IT neurons were assigned to one of the four genetically-defined subtypes based on laminae and gene expression, and to one of the projection subclusters based on projection patterns (Fig. S7B). Interestingly, subtypes that were further apart based on mean gene expression were also further apart based on clustering of projections (Fig. 6A), suggesting a relationship between gene expression and projection pattern within IT subtypes.

### The organization of projections across transcriptionally defined IT subtypes

Although the relationship between the IT1-IT4 subtypes defined by gene expression and the 19 subclusters defined by projection patterns was complex— projection patterns were largely mixed across the four IT subtypes—the large sample size enabled us to discern several novel relationships. We identified both qualitative and quantitative relationships between gene expression and projection pattern, including projection patterns specific to transcriptionally defined subtypes. These findings are detailed below.

Perhaps the most striking observation was the identification of a specific projection subcluster, which we denote ITi-Ctx, found almost exclusively in the two deep-layer subtypes IT3 and IT4, but absent in IT1 or IT2 (Fig. 7A). ITi-Ctx neurons are a subset (leaves 14 and 15 in Fig. 5C) of ITi neurons which project exclusively to the ipsilateral cortex, but not to the contralateral cortex or the striatum. Whereas ITi-Ctx neurons accounted for 13% (20 / 155) of IT3 neurons and 34% (45 / 123) of IT4 neurons, it accounted for only 2% (6 / 303) of IT1 neurons and fewer than 1% (1 / 145) of IT2 neurons (Fig. 7A; p < 1e-4 comparing IT3 to either IT1 or IT2, and p < 5e-15 comparing IT4 to either IT1 or IT2, all using Fisher’s exact test with Bonferroi correction). These ITi-Ctx neurons would have been difficult to detect using conventional retrograde tracing because they are defined by a combinatorial projection pattern, i.e. projection to ipsilateral but not contralateral cortex.

**Fig. 7.**
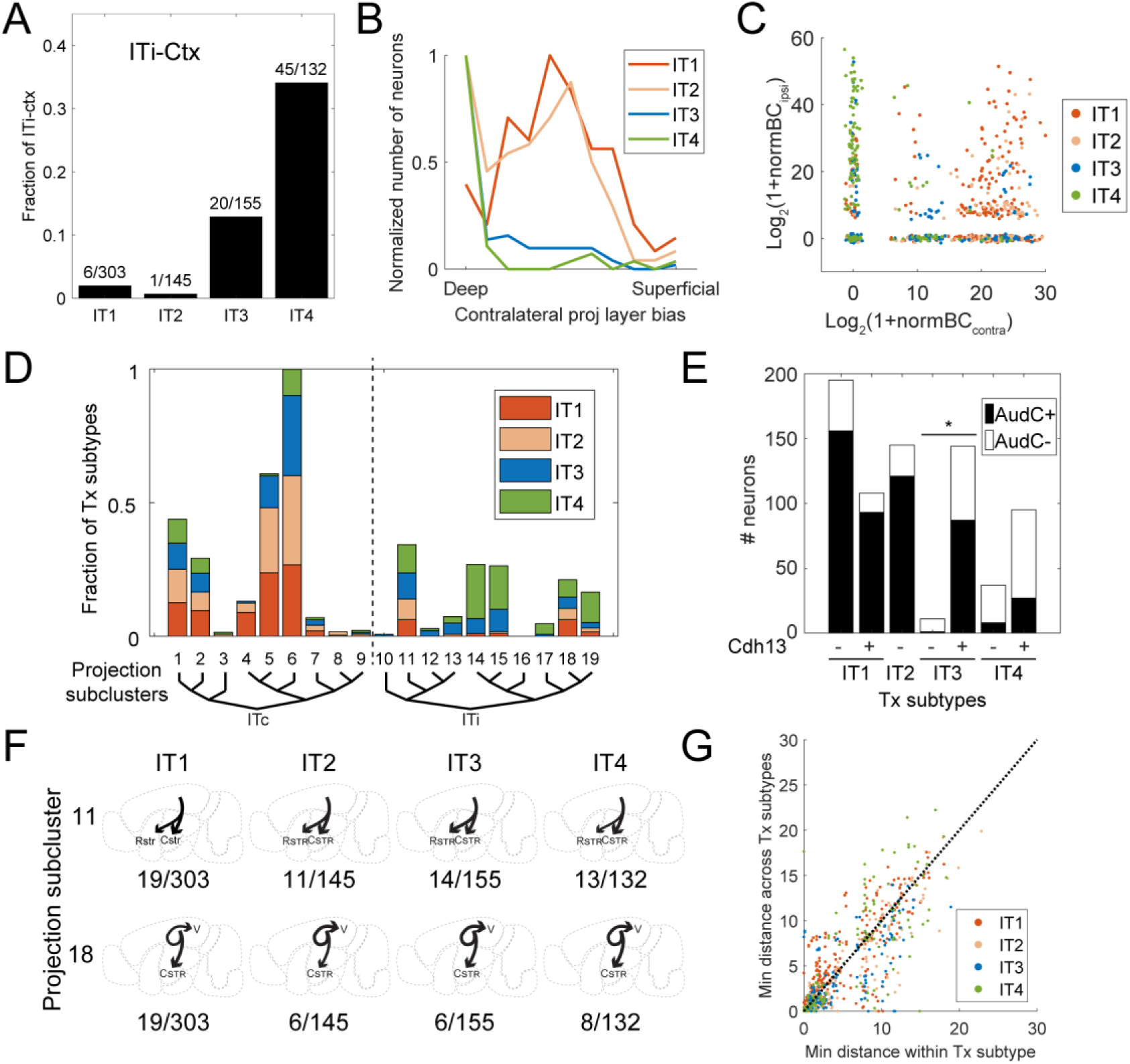
Projections across IT subtypes defined by gene expression. (A) The fraction of ITi-Ctx neurons in each indicated IT subtype defined by gene expression. The number of ITi-Ctx neurons in each subtype and the total numbers of each subtype are indicated on top of each bar. Inset: the mean projection pattern of ITi-Ctx neurons. (B) Histograms of the layer bias of contralateral projections of neurons in each IT subtype. The histograms are normalized so that the maximum value for a bin is 1 for each subtype. See STAR Methods for the definition of projection layer bias. (C) The Log normalized barcode count of projections to the contralateral auditory cortex (x-axis) is plotted against that of projections to the ipsilateral visual and somatosensory cortex (y-axis) for each neuron. Neurons are color-coded by IT subtypes defined by gene expression. (D) The fractions of neurons of IT subtypes defined by gene expression that belong to each indicated IT projection leaf subcluster. All bars belonging to a transcriptionally defined subtype sum to 1 across the whole plot. The projection cluster labels correspond to those in Fig. 5C. Vertical dashed line separates ITc and ITi subclusters. (E) The number of neurons with (AudC+) or without (AudC-) projections to the contralateral auditory cortex in each IT subtype defined by gene expression. Neurons in each IT subtype are further divided into those expressing Cdh13 and those that do not. *p < 0.005 using Fisher’s exact test after Bonferroni correction. (F) Two example projection leaf subclusters that were shared across all four IT subtypes defined by gene expression. Projection diagrams indicate example neurons from the indicated subtypes. The number of neurons belonging to a projection subcluster and a transcriptionally defined subtype and the total number of neurons in the transcriptionally defined subtype are indicated below each example. (G) The minimum distance in projections from a neuron to any neuron within the same subtype defined by gene expression (x-axis) or in a different subtype (y-axis). Neurons are color-coded by IT subtypes defined by gene expression. Dashed line indicates same minimum distance within and across subtypes. See also Fig. S7.

Second, we also found that when IT3 and IT4 neurons had contralateral projections, the projections specifically targeted the deep layers of the contralateral cortex (Fig. 7B; p < 5e-8 comparing IT3 to IT1 or IT2, and p < 5e-7 comparing IT4 to IT1 or IT2, all using Fisher’s exact test with Bonferroni correction). By contrast, the contralateral projections of IT1 and IT2 neurons terminate throughout all cortical layers. This result is consistent with, and expands upon, previous observations using classical anterograde tracing (Tasic et al., 2018) showing that a subset of IT3 neurons labeled by Penk-Cre project preferentially to deep layers.

Third, we found that if an IT3 or IT4 neuron had an ipsilateral projection, then it was unlikely to also have a contralateral projection. By contrast, neurons in IT1 and IT2 often projected both ipsi- and contralaterally (Fig. 7C; p < 5e-4 comparing IT3 to either IT1 or IT2, and p < 5e-16 comparing IT4 to either IT1 or IT2 using Fisher’s exact test with Bonferroni correction; see STAR methods). Because the presence of a contralateral projection is the characteristic that distinguishes the ITc from the ITi projection subcluster, most (84%; 377/448) IT1 or IT2 neurons were ITc, whereas fewer than half (43%; 123/287) of IT3 or IT4 neurons were ITc (Fig. 7D). These features of projections specific to IT3 and IT4 subtypes reinforce the observation that the rules governing the projections of IT3 and IT4 neurons are distinct from those governing IT1 and IT2 neurons.

Finally, we discovered finer quantitative differences across individual subtypes defined by gene expression. Although we identified two projection patterns (contralateral projections to deep layers and ITi-Ctx projections) shared by both IT3 and IT4, these two projection patterns were differentially enriched between the two subtypes. ITi-Ctx projections were both stronger (Fig. S7C, p = 0.012 comparing IT3 and IT4 using rank sum test) and more frequent in IT4 subtype compared to those in IT3 (Fig. 7D; 13% in IT3 compared to 34% in IT4 neurons, p = 3e-5 using Fisher’s exact test). In contrast, contralateral projections were twice as frequent in IT3 neurons compared to IT4 (Fig. 7D; 89 / 155 IT3 neurons compared to 34 / 132 IT4 neurons, p = 6e-8 using Fisher’s exact test). Within a subtype defined by gene expression, projections further correlate with gene expression: The probability of projecting contralaterally was higher among IT3 neurons with high Cdh13 expression (p = 0.001, Fisher’s exact test, Fig. 7E), an observation further confirmed using CTB retrograde tracing (Fig. S7D, p < 0.0001 using rank sum test).

Although the analyses above support the idea that gene expression and laminar position are correlated with projection pattern, the relationship between IT subtypes defined by gene expression and subclusters defined by projection patterns was in general complex (Fig. 7D). Thus, although some projection patterns were enriched in some subtypes [e.g., ITi-Ctx neurons (leaf 14 and 15) and ITc neurons of leaf 4 and 5; Fig. 7D], no projection subcluster was specific to a single subtype. Indeed, in many cases, the same projection subcluster could be found in all four IT subtypes; two examples of projection subclusters found in all four transcriptomic subtypes are shown in Fig. 7F. Such shared projection patterns across subtypes were common: 54% of IT neurons (400 / 735) were more similar in projections to a neuron of a different IT subtype than to any neuron of its own subtype (Fig. 7G; Fig. S7E-G). This mixed projection pattern is in contrast to the clear separation of projection patterns across the three higher-level classes of neurons (IT/PT-l/CT). Thus whereas at the top levels of the hierarchies, the classical partitioning into IT/PT-l/CT neurons captures the correlational structure of multiple cellular properties, including gene expression, laminar position and projection pattern, our results suggest that further subclustering based on one experimental property (such as gene expression) leads to categories that do not map neatly onto clustering based on another property (such as projections).

## Discussion

Here we have described BARseq, a barcoding-based neuroanatomical method that can relate high-throughput projection mapping with other neuronal properties at cellular resolution through *in situ* sequencing. As a proof of principle, we applied BARseq to the projections in the auditory cortex. BARseq of 3579 neurons recapitulated the organization of cortical projections into the classic IT/PT-l/CT subtypes. Within these classic subtypes, BARseq also revealed the impressive diversity of projection patterns: We observed 264 distinct projection patterns, falling into as many as 25 distinct clusters. We then combined BARseq with single cell analysis of gene expression of an additional 735 IT neurons. We identified projection patterns exclusive to, or enriched in, specific transcriptionally defined subtypes. BARseq thus revealed rich and complex relationships between gene expression and projection patterns that would have been difficult to uncover using conventional transcriptomic and/or anatomical techniques.

### Multiplexed projection mapping using cellular barcodes

BARseq achieves multiplexed projection mapping by matching barcode sequences found at the soma with sequences recovered at projection targets. This strategy, which differs fundamentally from that used by traditional single-neuron tracing experiments based on one or a few distinct tracers, confers both unique advantages and limitations upon BARseq.

BARseq fundamentally differs from conventional optical approaches in that projections are determined by matching barcodes, not tracing. Barcode matching, unlike optical tracing, does not accumulate error over distance. In conventional optical approaches to mapping projections, axons are reconstructed by observing continuity between successive optical or physical sections. Any lost or distorted section may result in an error, and in general the probability of error increases exponentially with the length of the axonal projection and the number of neurites multiplexed. For example, even with a low error rate of 1% per 50 µm, more than half of axons traced would be in error within 5 mm. By contrast, because BARseq relies on matching sequences to reconstruct projections, errors do not accumulate for long axons: A distant subcortical projection is just as reliably matched to its source as a projection to a nearby cortical area. Furthermore, the high diversity of barcodes (tens of millions in our experiments) can uniquely label tens of thousands of neurons in a single experiment. Therefore BARseq can determine projection patterns for orders of magnitude higher densities of neurons (hundreds to thousands of neurons per cortical area, and tens of thousands neurons per brain) than even the most advanced state-of-the-art multiplexed optical tracing methods [∼50 neurons per cortical area (Guo et al., 2017; Lin et al., 2018)], without the need of specialized high-speed microscope that is often required for advanced anatomical techniques.

In BARseq, the spatial resolution at which projections are resolved is determined by the size of the cubelets dissected. In this study, we chose to dissect brains relatively coarsely—only sufficient to distinguish among brain areas and between superficial and deep cortical layers, but this resolution was sufficient to resolve the organization of projections across neuronal subtypes. For questions requiring higher spatial resolution, BARseq can be further improved with laser capture microdissection (Huang et al., 2018) or direct *in situ* sequencing of projection barcodes, thus potentially resolving axonal projections and dendritic morphology at subcellular resolution. Such an approach would yield an “infinite color Brainbow,” allowing reconstruction of densely labeled neurons.

### The organization of projections across neuronal subtypes

The partitioning of neurons into types and subtypes is useful to the extent that these classes can infer multiple neuronal properties. The utility of these classes and subclasses arises from the fact that these properties co-vary. For example, knowing whether a cortical neuron is excitatory or inhibitory, or the excitatory subtype (IT/PT-l/CT) to which it belongs (Harris and Shepherd, 2015), allows us to infer a great deal about the neuron’s pattern of gene expression, morphology, connectivity and projection pattern. The partitioning of cortical neurons into excitatory vs. inhibitory, and of excitatory neurons into IT/PT-l/CT subtypes, is well supported by many lines of data.

Within these major subtypes of excitatory neuron, the predictive value of further subdivisions has been less clear. It has been reported that some PT-l subtypes defined by gene expression correspond to specific projection patterns (Economo et al., 2018), and some CT subtypes show subtype-dependent biases in their projections (Chevee et al., 2018). However, a systematic approach to determining the co-variance of two or more properties would require high-throughput measurements of multiple properties simultaneously, a feat that would be challenging using previous approaches. Indeed, it has only recently been possible to obtain such data on even a single property such as gene expression or projection pattern (Kebschull et al., 2016a).

The combination of BARseq with *in situ* transcriptional mapping has the potential to systematically determine the co-variation of multiple properties. As a proof-of-principle, we combined BARseq and FISH to explore the projections of transcriptionally defined IT subtypes. Using this approach, we identified a novel projection pattern (ITi-Ctx) restricted almost entirely to two transcriptionally defined IT subtypes (IT3 and IT4). We also identified several other ways in which patterns of gene expression covaried with projection patterns (Fig. 7). However, in marked contrast to the one-to-one relationship between top-level subtypes (IT/CT/PT-l) defined by gene expression and projection pattern, we did not observe a simple mapping of these properties within IT neurons.

The mixed projection patterns observed across IT subtypes defined by gene expression are unlikely to have resulted from errors in classifying neurons to subtypes, because we were able to identify both specific and promiscuous projection patterns in the same IT subtypes. This question can be further tested by correlating projections with more marker genes to better define IT subtypes by gene expression. To achieve this, BARseq can potentially be combined with highly-multiplexed FISH (Chen et al., 2015; Codeluppi et al., 2018; Eng et al., 2019; Shah et al., 2016) or *in situ* sequencing methods (Ke et al., 2013; Lee et al., 2014; Wang et al., 2018) to correlate the expression of dozens to hundreds of genes to projections. Such an approach could further dissect the relationship between projections and gene expression beyond IT subtypes, and has the potential to generate a comprehensive understanding of long-range projections and gene expression at cellular resolution.

### Rosetta Brain

Neuronal types differ by a combination of multiple properties (Cadwell et al., 2016; Economo et al., 2018; Paul et al., 2017), including anatomical characteristics such as morphology and connectivity (Economo et al., 2016; Gerfen et al., 2016), molecular characteristics such as gene (Luo et al., 2017; Tasic et al., 2016; Tasic et al., 2018; Zeisel et al., 2015) or protein expression, and functional characteristics such as behaviorally-evoked activity (Cadwell et al., 2016). Defining the biological functions of neuronal classes thus requires a multi-faceted approach that combines measurements of several neuronal properties together at cellular resolution (Huang and Paul, 2019; Zeng and Sanes, 2017). The ability to relate diverse characteristics of many single neurons simultaneously, in a co-registered fashion, within single brains, is an important challenge in neuroscience (Marblestone, 2014). Even simultaneous co-registration of two characteristics can be challenging, but the few successes in this arena have led to insights about the functional organization of neural circuits (Bock et al., 2011; Raj et al., 2018; Sorensen et al., 2015).

Co-registering multiple cellular properties at single-cell resolution is crucial for understanding biological systems with heterogeneous cell types, such as the nervous system. By sequencing barcodes *in situ*, BARseq offers a solution to incorporate high-throughput measurements based on cellular barcoding, such as lineage tracing (Raj et al., 2018), high-throughput screening (Feldman et al., 2018), and projection mapping (Kebschull et al., 2016a) into such integrated approaches. In this study, we focused on combining projections with adult gene expression and the laminar location of neuronal somata. Adapting BARseq for other barcoding techniques and adapting additional *in situ* methods, such as *in vivo* two-photon imaging, to combine with BARseq would yield a “Rosetta Brain”—an integrative dataset that could constrain theoretical efforts to bridge across levels of structure and function in the nervous system (Marblestone, 2014).

## Acknowledgements

The authors would like to acknowledge Kay Tye, Gordon Shephard, and Jessica Tollkuhn for useful discussions, and Barry Burbach, Nour El-Amine, and Stephen Hearn for technical support. This work was supported by the following funding sources: National Institutes of Health [5RO1NS073129 to A.M.Z., 5RO1DA036913 to A.M.Z.]; Brain Research Foundation [BRF-SIA-2014-03 to A.M.Z.]; IARPA MICrONS [D16PC0008 to A.M.Z.]; Simons Foundation [382793/SIMONS to A.M.Z.]; Paul Allen Distinguished Investigator Award [to A.M.Z.]; postdoctoral fellowship from the Simons Foundation to X.C. This work was performed with assistance from CSHL Shared Resources, which are funded, in part, by the Cancer Center Support Grant 5P30CA045508.

## Author contributions

X.C. and A.M.Z conceived the study. X.C., H.Z. and Y.S. optimized and performed BARseq. X.C. and Y.S. performed BARseq in combination with FISH and in Cre labeled mice. J.M.K, and H.Z. performed MAPseq. X.C. performed single-cell RNAseq. K.M. and Z.J.H generated Cre-labeled mice. X.C. and A.M.Z. analyzed the BARseq and MAPseq data. X.C., S.F., and J. G. analyzed the single-cell RNAseq data. X.C. and A.M.Z. wrote the paper.

## Materials & Correspondence

All correspondence and material requests should be made to A.M.Z

## Supplementary Figure and Table legends

**Fig. S1, related to** **Fig. 2** **and STAR Methods.** Optimization of *in situ* barcode sequencing for brain slices. (A) Amplification of barcodes in brain slices in the indicated reaction chambers. Scale bars = 100 µm. (B) Merged images of rolonies (yellow) generated in barcoded brain slices and the residual GFP signals (cyan) with the indicated time of pepsin treatment. Scale bars = 100 µm. (C) Comparison of barcode amplicons generated using BaristaSeq (a), the original padlock method (b), and FISSEQ (c). Scale bars = 50 µm. (D) Sequencing images of cycles 2, 4 and 6 of barcoded brain slices sequenced using SOLiD sequencing chemistry (*top*) and using Illumina sequencing chemistry (*bottom*). Imaging conditions were kept constant throughout each sequencing run. Scale bars = 100 µm. (E) Average signal-to-noise ratio of Illumina (red) and SOLiD (blue) sequencing *in situ* over cycles. Error bars indicate the standard errors for the SNR for pixels. (F-H) Sequencing quality and signal intensity of individual base calls (F), mean signal intensity over cycles (G), and the fraction of the bases over cycles (H) are plotted. (I) The distribution of barcode counts in contralateral auditory cortex compared to those in the olfactory bulb (negative control). Dashed line indicates threshold above which a projection is considered real.

**Fig. S2, related to** **Fig. 3**. Comparison of gene expression in barcoded neurons to that in non-barcoded neurons. (A) Mean log normalized expression of each gene averaged over all barcoded (x-axis) and non-barcoded (y-axis) neurons. The gene expression is regressed linearly using the percentage of mitochondrial genes. The diagonal line indicates equal expression in barcoded vs. non-barcoded cells. (B) Fraction of variance explained by the same top principal components (x-axis) in non-barcoded neurons (blue), barcoded neurons (red), and randomized non-barcoded neurons (yellow). Vertical bars along the non-barcoded line indicate 95% confidence interval. (C) Heat maps of the expression of top genes in the first six principal components from non-barcoded neuros.

**Fig. S3, related to** **Fig. 4** **and STAR Methods.** Locations of neurons in the auditory cortex mapped by BARseq. (A) Representative low-resolution image of a brain slice showing barcoded GFP+ cells (cyan) and DAPI staining in a typical injection site in the auditory cortex for BARseq and MAPseq experiments. This image was not acquired in any BARseq or MAPseq brains because the workflow of these experiments does not preserve whole brain slices. Scale bar = 500 µm. (B)(C) Representative images of FISH against Cux2 (B) and Fezf2 (C) in two adjacent slices. Scale bars = 50 µm. (D) Normalized layer boundaries determined in three pairs of slices across the auditory cortex. In each slice, the thickness of the cortex was normalized to that in the BARseq brains, and the boundary positions were scaled accordingly. (E) Violin plots of the laminar distribution of all BARseq neurons in XC9 and XC28 (All) and those with (Proj) or without (Non-proj) detected projections. Individual neurons (red) are plotted on top of smoothed distribution plots.

**Fig. S4, related to** **Fig. 5**. Projections of the auditory cortex revealed by BARseq. (A) Triple retrograde tracing of neurons projecting to the rostral striatum (CTB-647), the caudal striatum (CTB-488), and the tectum (RetroBeads). (B) A representative image of triple retrograde labeling in the auditory cortex showing neurons projecting to the rostral striatum (magenta), the caudal striatum (cyan), and the tectum (yellow). Scale bar = 100 µm. (C) Venn diagram showing the number of neurons projecting to each of the three areas. (D) The strengths of the indicated projection (y-axes) of individual neurons of the indicated classes are plotted against their laminar position (x-axes). The mean projection strengths are marked by black lines. The layer boundaries are marked by dotted vertical lines.

**Fig. S5, related to** **Fig. 5** **and STAR Methods.** Hierarchical clustering of projection neurons in the mouse auditory cortex. (A) Positive (green) or negative (red) Spearman correlation coefficients among projections to the indicated areas in the indicated classes. Only correlations that were statistically significant after Bonferroni correction were shown. (B) The workflow of the hierarchical clustering. (C) Comparison of the original projection strengths (blue) and the filtered projection strengths (red) for two example neurons. (D) The fraction of variance explained (y-axis) using non-negative matrix factorization (NMF; blue), individual projections (red), and PCA (black). (E) The fractions of neurons that remain in the same class-level clusters (y-axis) when filtering the projection data with the indicated number of projection modules (x-axis) compared to the clusters without filtering. (F) Comparison of clusters obtained using k-means (*upper row*), spectral clustering (*middle row*), and Louvain community detection (*lower row*) at the indicated hierarchies. All clusters are color-coded onto the same t-SNE plot. The colors are randomly assigned to individual clusters. (G) The distribution of the maximum cluster probability for individual neurons when classified using all 11 projection areas (a) or 10 projection areas (b-l). For classification using 10 projection areas, the unused projection area is labeled on top of each graph. (H) The fraction of well-classified neurons in projection leaf subclusters. The subcluster labels correspond to those in Fig. 5C and the major classes they belong to are labeled below.

**Fig. S6, related to** **Fig. 5**. Consistent projections and laminar distribution across brains. (A) The fractions of neurons from each brain that belong to each indicated high-level projection subcluster. All bars belonging to a brain sum to 1 across the whole plot. (B) Differences in normalized entropy (x-axis) of individual subclusters between the two brains are plotted against the negative logarithm of the p values (y-axis). The subclusters are color-coded according to their class-level divisions as indicated. The p values are shown without multiple testing correction. The red vertical dashed line indicates no difference in entropy, and the black horizontal dashed line indicates significance level after Bonferroni correction. (C) Differences in mean laminar locations (x-axis) of individual subclusters between the two brains are plotted against the negative logarithm of the p values (y-axis). The subclusters are color-coded according to their class-level divisions as indicated. The p values are shown without multiple testing correction. The red vertical dashed line indicates no difference in the mean laminar locations, and the black horizontal dashed line indicates significance level after Bonferroni correction.

**Fig. S7, related to** **Fig. 6** **and** **Fig. 7**. Projections of IT subtypes defined by gene expression. (A) t-SNE plots of projections color-coded by brain indices. (B) t-SNE plots of projections of neurons of the indicated IT subtypes defined by gene expression. Neurons of the indicated IT subtypes are color-coded by high-level projection subclusters, and other neurons are grayed out. (C) Histograms of log normalized barcode counts of projections to the ipsilateral visual cortex in IT3 and IT4 neurons. (D) Cumulative probability distribution of Cdh13 expression in contralateral projection neurons and ipsilateral projection neurons as determined by FISH and CTB retrograde tracing. (E) The minimum distance from a neuron to any neuron within the same projection subcluster (x-axis) or in a different projection subcluster (y-axis). Neurons are color-coded by high-level projection subclusters. Dashed line indicate same minimum distance within and across subtypes. (F) For each neuron belonging to an indicated IT subtype defined by gene expression, the number of IT subtypes containing neurons with the same binary projection pattern was counted (possible values range from 1 to 4 subtypes). The histogram of this value was then plotted. (G) Histogram of the number of transcriptionally defined subtypes a binary projection pattern was found in.

**Table S1, related to** **Fig. 2**. Comparison between BARseq and CTB retrograde tracing. Each row represents a single neuron recovered from BARseq with visible GFP signal from the barcodes and good sequencing quality (quality score > 0.75, column 4). The first four columns indicate the raw barcode counts in the olfactory bulb (OB), contralateral auditory cortex (c1), the cortical area surrounding the contralateral auditory cortex with CTB signals visible to the naked eyes (c2), and an even larger cortical area surrounding the tracer area with CTB signals visible under the microscope (c3). The rest of the columns indicate minimal sequencing scores and whether the cell projects contralaterally based on CTB and/or BARseq.

**Table S2, related to** **Fig. 3**. Correlation of gene expression and projections using BARseq. Each row represents a single barcoded neuron with a minimum quality score over 0.8. Columns 2-7 indicate barcode molecule counts in each collected area, and column 8 indicates total barcode molecule counts. Columns 9 and 10 indicate whether the cells expressed Slc17a7 and/or Gad2.

**Table S3, related to** **Fig. 4**. Summary of BARseq datasets. The number of target areas collected, the number of slices sequenced in situ, The number of barcodes sequenced per brain from the projection sites, the number of cells sequenced per brain from the auditory cortex, and the number of *in situ* sequenced barcodes matching barcodes at the projection sites with or without quality filtering are indicated for each brain. *In XC14, the auditory cortex was sequenced *in vitro*, and no further filtering was applied in addition to those during standard MAPseq data processing. **In experiments combining BARseq and FISH, low-quality neurons were filtered out before mapping to projection pattern. The number of cells in ACx were thus those after filtering. *** In the experiments combining BARseq and Cre labeling, the quality filtering was done similar to the experiments combining BARseq and FISH. The number of cells in ACx listed thus only included high-quality neurons. The number of filtered cells here indicate projection neurons double labeled by the Cre.

## STAR Methods

### Viruses, constructs, and oligos

The plasmid encoding the Sindbis barcode library (JK100L2, https://benchling.com/s/EKtQttOe) is available from Addgene (#79785). The RT primer (XC1215), the padlock probe / sequencing primer (XC1164), and the fluorescent probe for visualization (XC92) were described previously (Chen et al., 2018).

For validation of barcode sequencing in brain slices, we used a barcode library previously described by Kebschull et al. (2016a). The library contained 1.5 million known 30-nt random barcode sequences, which represented ∼97% of all barcodes in the library. For BARseq experiments, we used a separate diverse barcode library with ∼10^7^ diversity (Han et al., 2018). This library was not fully sequenced *in vitro*.

### Animals and tissue processing

All animal procedures were carried out in accordance with Institutional Animal Care and Use Committee protocol 16-13-10-07-03-00-4 at Cold Spring Harbor Laboratory. Eight to nine week old male C57BL/6 mice were injected in the left auditory cortex at -4.3 mm ML, -2.6 mm AP from bregma, with 140nL 1:3 diluted Sindbis virus at each of the following depths (200 µm, 400 µm, 600 µm, and 800 µm) at 30° angle. For samples prepared for BaristaSeq only, we transcardially perfused the animal with 10% formalin, then postfixed the tissue for 24 hrs. We then cryo-protected the brain in PBS with 10% sucrose for 12 hrs, 20% sucrose PBS for 12 hrs, and 30% sucrose PBS for 12 hrs. We then embedded the brain in OCT (Electron Microscopy Sciences) and cryo-sectioned to 14 µm slices onto SuperFrost Plus slides (VWR).

For BARseq samples, we trans-cardially perfused the animal with PBS 43-45 hrs post-injection. We cut out the left auditory cortex from the brain and post-fixed it in 10% formalin at 4 °C for 8 hrs, and snap-froze the rest of the brain on a razor blade on dry ice. The snap-frozen brain (without the injection site) was then processed for conventional MAPseq as described (Kebschull et al., 2016a). The post-fixed auditory cortex was cryo-protected, embedded, and cryo-sectioned as described above.

For combined BARseq/FISH experiments in Fig. 3D-F, we injected 8-week old C57BL/6 male mouse at -4 mm ML, -2.6 mm AP from bregma, with 140nL 1:3 diluted Sindbis virus at depths 200 µm, 400 µm, 600 µm, and 800 µm straight down. After 24 hours, the animal was processed as described above for BARseq experiments. For the experiment in Fig. 6D-E and Fig. 7, 8-week old C57BL/6 male mice were injected at -4.3 mm ML, -2.6 mm AP from bregma with 140nL 1:3 diluted Sindbis virus at depths 300 µm, 600 µm, and 900 µm at 30° angle. After 24 hours, the animal was processed as described above for BARseq experiments, except that the injection site was flash frozen in OCT using isopentane and liquid nitrogen immediately after dissection from the brain. These brains were cryo-sectioned to 14 µm sections using a home-made tape-transfer system (Pinskiy et al., 2015) and glued to slides using NOA 81 (Norland) to reduce tissue distortion.

To compare BARseq to retrograde tracing, we injected 140 nL Alexa 647 labeled cholera toxin subunit B (CTB) into the right auditory cortex at 4.3 mm ML, -2.6 mm AP at multiple depths (200 µm, 400 µm, 600 µm, and 800 µm) at 30° angle. After 48 hrs, we injected 140 nL 1:3 diluted JK100L2 virus into the left auditory cortex at -4.3 mm ML, -2.6 mm AP, with 140nL Sindbis virus at each depth (200 µm, 400 µm, 600 µm, and 800 µm) at 30° angle. After another 44 hrs, the animals were then processed as for conventional BARseq samples.

For single-cell RNAseq comparing barcoded cells to non-barcoded cells, 8-week old C57BL/6 male mice were injected at -4.3 mm ML, -2.6 mm AP from bregma with 140nL 1:3 diluted Sindbis virus at depths 300 µm, 600 µm, and 900 µm at 30° angle. After 24 hours, the animal was processed for single-cell dissociation as described below.

### BaristaSeq

BaristaSeq on cultured neurons was performed as described (Chen et al., 2018). Briefly, the neurons were fixed in 10% formalin, washed in PBST (PBS with 0.5% tween-20), and dehydrated in 70%, 85%, and 100% ethanol for an hour. After rehydration in PBST, we incubated the samples in 0.1M HCl for 5 mins, followed by three PBST washes. We then reverse transcribed the samples [1 U/µl RiboLock RNase inhibitor (Thermo Fisher Scientific), 0.2 µg/µl BSA, 500 µM dNTPs (Thermo Fisher Scientific), 1 µM RT primer, and 20 U/µl RevertAid H Minus M-MuLV reverse transcriptase (Thermo Fisher Scientific) in 1× RT buffer] at 37 °C overnight. After reverse transcription, we crosslinked the cDNAs in 50 mM BS(PEG)_9_ for 1 hr and neutralized with 1M Tris-HCl for 30 mins. We then gap-filled and ligated padlock probes [100 nM padlock probe, 50 µM dNTPs, 1 U/µl RiboLock RNase inhibitor, 20% formamide (Thermo Fisher Scientific), 0.5 U/µl Ampligase (Epicentre), 0.4 U/µl RNase H (Enzymatics), and 0.2 U/µl Phusion DNA polymerase (Thermo Fisher Scientific) in 1× ampligase buffer supplemented with additional 50 mM KCl] for 30 mins at 37 °C and 45 mins at 45 °C. Following PBST washes, we performed rolling circle amplification (RCA) [20 µM aadUTP, 0.2 µg/µl BSA, 250 µM dNTPs, and 1 U/µl ϕ29 DNA polymerase in 1× ϕ29 DNA polymerase buffer supplemented with 5 % additional glycerol] overnight at room temperature. After crosslinking the rolonies using BS(PEG)_9_ and neutralizing with Tris-HCl, we hybridized 2.5 µM sequencing primers or 0.5 µM fluorescent probes in 2× SSC with 10 % formamide, washed three times in the same buffer, and proceeded to sequencing or imaging.

For BaristaSeq on brain tissues, we tested three commercially available reaction chambers that were physically compatible with our samples (Fig. S1Aa-c), and found that the HybriWell-FL sealing system was the only system that did not inhibit rolony formation (Fig. S1Ab). The ImmEdge hydrophobic barrier pen also produced good amplification (Fig. S1Ad), but the HybriWell-FL system offered better control of liquid evaporation during heating steps and easier handling. All slides with brain slices were thus first sealed in HybriWell-FL chambers (22 mm x 22 mm x 0.25 mm; Grace Bio-labs) for reactions. The brain slices were washed three times in PBS supplemented with 0.5% Tween-20 (PBST), followed by a pepsin digestion step. This step was necessary to increase accessibility of fixed RNAs (Fig. S1B) and to reduce the GFP signal from the cells (cyan in Fig. S1B), which may interfere with sequencing signals. We found that 3 mins of 0.2% pepsin digestion in 0.1 M HCl at room temperature greatly increased rolony formation (Fig. S1Bb) compared to no pepsin treatment (Fig. S1Ba), whereas 5 mins of pepsin digestion caused excessive tissue loss (Fig. S1Bc). We therefore used 3 mins of pepsin digestion for BaristaSeq in most brain slices, but the optimal timing could vary with different fixation conditions. We then proceeded with ethanol dehydration, followed by reverse transcription, padlock gap-filling and ligation, and RCA as described above for cultured neurons.

To sequence the barcodes using Illumina sequencing chemistry, we based our sequencing protocol on the HiSeq recipe files (Chen et al., 2018) and reduced the CRM incubation time to two minutes each due to efficient heat transfer of the reaction chambers. We also increased the PBST washes to four to eight times after the IRM reactions to counteract the increased background staining in tissue slices. For slices sectioned using the tape-transfer system (including XC54, XC75, XC91, and XC92), we further incubated the samples in 0.4% MMTS (Pierce) in PBST at 60 °C for 3 mins after CRM and SB3 wash. This optimized Illumina sequencing protocol resulted in an average signal-to-noise ratio 39 ± 4 in the first six sequencing cycles (Fig. S1D, E), ∼10-fold higher than that of sequencing by ligation (SOLiD; 4 ± 1) used by other sequencing methods (Ke et al., 2013; Lee et al., 2014).

### Imaging

All imaging except for the experiments in Fig. 2C-F and Fig. 3A-C was performed on an UltraView VoX spinning disk confocal microscope (Perkin Elmer) with Volocity 6.3 software as previously described (Chen et al., 2018). The sequencing channels were: Channel G, 514 nm laser excitation, 405/440/514/640 quad dichroic, 550/49 emission filter, exposure time 500 ms; Channel Y, 561 nm laser excitation, 405/488/561/640 dichroic, 445/60 and 615/70 dual band emission filter, exposure time 120 ms; Channel A, 640 nm laser excitation, 405/488/561/640 dichroic, 676/29 emission filter, exposure time 250 ms; Channel C, 640 nm laser excitation, 405/488/561/640 dichroic, 775/140 emission filter, exposure time 500 ms. The experiments in Fig. 2C-F were produced on a Zeiss LSM 710 laser scanning confocal microscope with Zeiss Zen software as previously described (Chen et al., 2018). The experiment in Fig. 3A-C was imaged on a Nikon TE2000 with Crest X-light spinning disk, Photometrics Prime 95B camera, and an 89north 7-line LDI laser. The scope was controlled by micro-manager (Edelstein et al., 2014). The four sequencing channels were imaged using the following settings: Channel G, excitation 520 nm laser, ZT443-518rpc-UF2 dual-band dichroic (Chroma), FF01-565/24-25 emission filter (Semrock), exposure time 600 ms; Channel T, excitation 555 nm laser, zt402/468/555/640rpc-uf2 quad-band dichroic (Chroma), FF01-585/11-25 emission filter (Semrock), exposure time 200 ms; Channel A, excitation 640 nm laser, zt402/468/555/640rpc-uf2 quad-band dichroic (Chroma), FF01-676/29-25 emission filter (Semrock), exposure time 300 ms; Channel C, excitation 640 nm laser, zt402/468/555/640rpc-uf2 quad-band dichroic (Chroma), FF01-725/40-25 emission filter (Semrock), exposure time 500 ms. The channels were calibrated as described previously (Chen et al., 2018).

For XC9 and XC28, a 3×1 stitched image at 10× was acquired for each slice, with each tile position including 11 z-positions with 10 µm step size. For XC75 and XC91, a 3×3 stitched image at 10× was acquired for each slice. Imaging time was about one minute per slice for XC9 and XC28, and three minutes per slice for XC75 and XC91. During each sequencing run, 24-36 slices on three slides were imaged together, resulting in a per-cycle imaging time of 0.5-1 hour. For XC75 and XC91, an additional 5×5 tiles of 20× image was acquired for the first sequencing cycle, with each tile position including 17 z-positions with 3 µm step size. This high-resolution imaging typically took 15 minutes per slice.

All images in figures were maximum projections of z-stacks shown after rolling ball background subtraction except for images in Fig. S1D, which were shown with only the camera blackpoint subtracted from the images.

### Base-calling

Max projection images were first processed through a median filter and a rolling ball background subtraction. The processed images were then corrected for channel bleed through. The barcodes were then base-called by picking the channel with the strongest signal. The sequencing quality score is defined as the intensity of the called channel divided by the root sum square of all four channels.

### Comparison to other *in situ* sequencing techniques

The original padlock probe-based barcode amplification was performed similar to BaristaSeq amplification as described above, except that the Stoffel fragment (DNA Gdansk) was used in place of Phusion DNA polymerase and the cDNA crosslink was done using 4% paraformaldehyde in PBST for 10 mins (Ke et al., 2013).

To perform targeted FISSEQ in brain slices, we processed the sample in the same way as in BaristaSeq in brain slices to the cDNA crosslink step. After cDNA crosslinking, we digested the RNAs [10 µl RiboShredder (Epicentre) and 5 µl RNase H in 1× RNase H buffer] for 1 hour at 37 °C. After washing the samples twice in water, we circularized the cDNAs [0.5 mM DTT, 1M Betaine, 2.5 mM MnCl_2_, and 1 U/µl Circligase II in 1× Circligase buffer] for 1 hour at 60 °C. After washing the samples with PBST, we hybridized 1.5 µM RCA primers in 2× SSC with 10% formamide for 1 hour. We then washed the samples three times with the same buffer, twice more with PBST, and proceeded to RCA as in BaristaSeq.

To compare Illumina sequencing chemistry to SOLiD sequencing chemistry for *in situ* sequencing in tissues, we performed SOLiD sequencing as described previously (Chen et al., 2018; Lee et al., 2014). To calculate the signal to noise ratio, we first converted the four-channel images into one single channel, taking the maximum value of the four channels for each pixel. We selected areas containing barcoded cells or areas containing tissues but no barcoded cells by thresholding as the signal and background areas. We then subtracted the black point of the camera from both the signals and the backgrounds and calculated the SNR. The SNR was calculated using the same selected areas in all six cycles.

### BARseq

Animals were injected and processed as described above. Cryo-sectioned brain slices were first imaged to generate DIC and GFP images, before they were processed for BaristaSeq. The GFP images from neighboring slices were aligned to each other manually to reduce deformation during sectioning. The sequencing images were then registered back to the GFP images to locate the positions of the neurons within the slice. Each basecall ROI was thus registered back to the aligned GFP images.

We dissected 12 projection sites from the frozen brains for sequencing. These 12 sites included the olfactory bulb, the orbitofrontal cortex, the motor cortex, the rostral and caudal striatum, the somatosensory cortex, the ipsilateral and contralateral visual cortex, the contralateral auditory cortex, the thalamus, and the tectum. The auditory cortex dissections also included the neighboring temporal association area, and the visual cortex collected included the posterior parietal cortex. Slice images, dissected areas, and their correspondence to the Allen Reference Atlas are available at Mendeley (see Data Availability). The projection sites were sequenced as described for MAPseq (Kebschull et al., 2016a).

For experiments using combination of BARseq and FISH (XC75 and XC91), each of the cortical projection sites was separated into upper and lower layers and collected separately, resulting in 18 projection sites.

### BARseq data processing

We first filtered the MAPseq generated barcodes so that all barcodes had at least 10 molecules but no more than 10000 molecules at the strongest projection site. We recovered 26840 barcodes using these criteria from the three brains (Table S3). We then matched these barcode sequences at the projection sites to those at the injection site, allowing three mismatches for conventional MAPseq or one mismatch for BaristaSeq. In the conventional MAPseq brain (XC14), 5082 out of 8418 barcodes were confirmed to be from the auditory cortex with more than 20 molecules per barcode at the injection site and were used for the subsequent analyses.

In BARseq experiments, the injection sites were sequenced *in situ* to 15 bases. The 15 bases read length *in situ* was sufficient to distinguish unambiguously all infected barcodes allowing one mismatch. For the XC9 brain, barcodes recovered through MAPseq had a mean hamming distance of 4.5 ± 0.7 (mean ± stdev; Fig. 5A). Only one pair (0.04%) out of 4841 barcodes had a hamming distance of 1 and 10 pairs (0.4%) out of 4841 had a hamming distance of 2. Because the sequencing experiment in Fig. 2 showed only a single error for 51 barcodes, each sequenced 25 bases, our sequencing error rate was approximately 1⁄(51 × 25) = 0.08%. Therefore, assuming that sequencing errors have no bias toward a particular base, the probability of matching an *in situ* barcode to the wrong MAPseq barcode, while allowing one mismatch, is 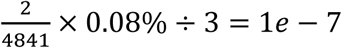. The probability of an *in situ* barcode matching to two MAPseq barcodes is 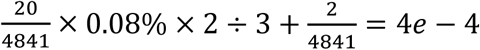. Although we cannot detect false positive matches, an ambiguous match could be detected. In the XC9 data, however, no ambiguous match between the *in situ* barcodes and the MAPseq barcodes has occurred.

In addition, XC9 had three pairs of barcodes whose first 15 bases were the same. These appeared to have arisen from amplification errors in homopolymer stretches of the same barcode rather than different barcodes, because each pair had a single in-del and had almost identical projection patterns. These three pairs were not recovered *in situ* and thus did not affect the analyses.

Similarly, out of 13581 total sequences, XC28 had 5 pairs of barcodes within one mismatch and 106 pairs of barcodes within two mismatches for the first 15 bases. No XC28 barcodes had identical sequences in the first 15 bases. The probability of a wrong match in XC28 is 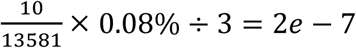, and the probability of an ambiguous match in XC28 is 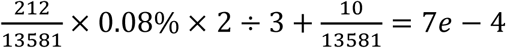. No actual ambiguous match was seen in XC28. Therefore, allowing one mismatch for a 15-base sequence was sufficient to match barcodes in the somata to those at the projection sites unambiguously for both brains.

In the two BARseq brains (XC9 and XC28), we sequenced 3237 cells *in situ*. Of all sequenced cells, 1806 (56%) cells had corresponding sequences at any projection site. The remaining cells had either low read qualities (possibly from having more than one barcode in the cell) or did not project to the examined areas (e.g., local interneurons, excitatory neurons that project to secondary auditory areas, and non-neuronal cells). We further filtered out barcodes with fewer than 10 molecules in the maximum projection area, removed neurons below the bottom of the cortex (these are likely persistent subplate neurons in the callosal commissure) and neurons in highly distorted slices (as judged by an abnormal cortical thickness). After filtering, 1309 neurons were used in the analyses.

### Comparison of BARseq to retrograde tracer

The animals were injected and processed as described above. We collected four target sites, including the olfactory bulb (negative control), the contralateral auditory cortex (i.e. the center of the CTB injection), the remaining areas where CTB was visible to the naked eye, and the surrounding areas where CTB was visible under a fluorescent microscope. The last three samples thus formed concentric rings around the CTB injection site. All three samples gave consistent results regarding contralateral projections (Table S1)

Before library preparation for BaristaSeq, we also imaged the Alexa 647 channel to locate retrograde-labeled neurons. These images were aligned to the sequencing images. We then only counted neurons clearly expressing GFP, which was essential to properly judge colocalization with CTB. To find the fraction of CTB labeled neurons that were also labeled by BARseq, we identified neurons labeled by both CTB and barcodes with a minimum sequencing quality of 0.75, and counted the number of neurons with barcodes in the contralateral auditory cortex above the noise floor. The noise floor was set to be the maximum count of individual barcodes recovered in the olfactory bulb.

### BARseq in Fezf2-2A-CreER::Rosa-CAG-loxP-STOP-loxP-tdTomato (Ai14) animals

Fezf2-2A-CreER::Rosa-CAG-loxP-STOP-loxP-tdTomato (Ai14) animals were generated and Cre recombination was induced at 6 and 7 postnatal weeks by intraperitoneal injection of Tamoxifen (Sigma-Aldrich T-5648) (100 mg/kg for each dose). A Fezf2-2A-CreER::Ai14 animal was injected at three postnatal months with barcodes in the auditory cortex as described above for BARseq. After 24 hours, the animal was anesthetized with isoflurane and decapitated. The injection site was punched out using a 2 mm diameter biopsy punch, and the rest of the brain was flash frozen for dissection of projection areas. The injection site punch was then post-fixed in 4% PFA for 24 hours, cryo-protected in 10%, 20%, and 30% sucrose in PBS for 12 hours each, mounted in OCT, and frozen. The punch was then cryo-sectioned to 14 um slices using the tape-transfer system and imaged for GFP, RFP, and DIC channels on a Nikon TE2000 microscope with a Crest X-light spinning disk confocal. The slices were then processed for BARseq as described above. Sequencing was done as described above for 14 cycles, except that the imaging was done on a Nikon TE2000 microscope with a Crest X-light spinning disk confocal. In addition, a set of sequencing images was taken after each cleavage step. This background image records the residual tdTomato signals in the sequencing images. After registering and subtracting the background image from sequencing images, we then median filtered the images, performed background subtraction, corrected for channel bleed-through, and registered sequencing images as described above. We also registered the “T” channel of the first sequencing cycle to the tdTomato images taken before library preparation. We then picked cell bodies from the sequencing images using the “Find Maxima” function in ImageJ, base-called for barcode sequences, and read out the intensity from the tdTomato images. We filtered out cells with sequencing quality lower than 0.7. We considered cells with tdTomato signal over 20000 as Fezf2+ neurons. These cells were visually inspected to make sure that they had good morphology and were labeled by tdTomato. These cells were then used for analyses.

### Validation of BARseq identified projections using retrograde tracing

To validate the striatal projections of PT-l and IT neurons, we injected red RetroBeads (LumaFluor) diluted 1:1 in PBS in the superior colliculus at -4.8 mm AP, -0.7 mm ML from bregma at depths 500 µm, 700 µm, 900 µm, 1100 µm, and 1300 µm (70 µl per depth) from the surface of the brain, Alexa 488 labeled CTB in the caudal striatum at -1.6 mm AP, -3.2 mm ML at depths 2.5 mm and 3 mm (50 µl per depth) from the surface of the brain, and Alexa 647 labeled CTB in the rostral striatum at 0.6 mm AP, -2 mm ML at depths 2.5 mm and 3 mm (50 µl per depth) from the surface of the brain. After 96 hrs, we perfused the animal and sliced the auditory cortex coronally into 70 µm slices. We then imaged the slices on an UltraView VoX spinning disk confocal microscope (Perkin Elmer).

### Validation of combination of BARseq and FISH

To correlate projections to the expression of Slc17a7 and Gad2, we collected the olfactory bulb, ipsilateral cortical areas (mainly the visual cortex), contralateral cortical areas (the auditory and visual cortex), the striatum, the thalamus, and the tectum. These areas were processed for MAPseq as described above and sequenced on an Illumina MiSeq. The injection site was cut out from the brain, post-fixed and cryo-protected for cryo-sectioning.

We cryo-sectioned the injection site to 14 µm slices using the tape transfer system. The brain slices were then processed for BARseq to the first cross-link step after reverse transcription. After neutralization of additional crosslinkers with Tris, we proceeded with ViewRNA ISH (Thermo Fisher Scientific) using a class 1 probeset for Slc17a7 and a class 6 probeset for Gad2 according to the manufacturer’s protocol. We then imaged each area around the injection site (identified by the presence of GFP) using a 20× 0.75NA objective on a spinning disk confocal for both FISH channels (RFP and Cy5), the GFP channel, and the DIC channel. We then stripped away the FISH probes using 80% formamide for 5 mins twice, followed by three washes in 10% formamide in 2× SSC, and two washes with PBST. We then proceeded to the padlock ligation step in BARseq and produced rolonies as described above.

The sequencing of rolonies was done as described above except that the first sequencing cycle was imaged first using a 20× 0.75NA objective without binning for all four sequencing channels and the DIC channel. This was followed by imaging using a 10× 0.45NA objective with 2× binning. All subsequent sequencing cycles were imaged at the lower resolution to increase throughput.

We then registered the high-resolution FISH images to the high-resolution first cycle sequencing images using the DIC channel. To match the high-resolution first cycle sequencing images to the low-resolution sequencing images, we down-sampled the first-cycle sequencing images by four folds and applied a Gaussian filter to mimic the lower optical resolution. The two images were then roughly aligned manually. We then extracted cell-body locations using the “Find Maxima” function in ImageJ, and further aligned the cell-body locations using Iterative Closest Points (Besl and McKay, 1992) in MATLAB (https://www.mathworks.com/matlabcentral/fileexchange/27804-iterative-closest-point). The barcodes were called from the low-res images using the cell-body locations extracted in the previous step, and the expression of Slc17a7 and Gad2 were determined manually by examining the overlap between FISH signals and barcode rolonies. We then filtered out cells with a minimum sequencing quality score of less than 0.8, and further removed debris based on morphology and homopolymer barcode sequences. The remaining 99 cells were analyzed for projection pattern and gene expression.

Of the 99 barcoded cells with high sequencing quality and good morphology (Fig. 3F; Table S2), 80 were excitatory neurons (Slc17a7+ and Gad2-), 3 were inhibitory neurons (Slc17a7- and Gad2+), and 16 were non-neuronal cells (Slc17a7- and Gad2-). The ratio between inhibitory and excitatory neurons was not significantly different from previous estimates [∼11.5% (Meyer et al., 2011), p = 0.13 using Fisher’s exact test]. Out of these 99 cells, 54% (54/99) projected to at least one of the sampled areas with strengths above the noise floor defined by the olfactory bulb (Table S2). Consistent with the fact that most projection neurons in the cortex are excitatory, all 54 projection neurons identified by BARseq expressed Slc17a7. The non-projecting excitatory neurons likely projected locally or to nearby cortical areas we did not sample.

### Estimation of laminar boundaries using fluorescent *in situ* hybridization (FISH)

To estimate the boundaries of cortical layers, we performed FISH against two known layer-specific marker genes, Cux2 (Custo Greig et al., 2013) (Fig. S3B), and Fezf2 (Fig. S3C). Cux2 was strongly expressed in L2/3 and only sporadically in other layers; Fezf2 was strongly expressed in L5 and weakly in L6. Because L4 is poorly defined in the auditory cortex (Linden and Schreiner, 2003), we omitted L4 and defined only the remaining two borders. We defined the L2/3 and L5 border as below the Cux2 band and above the strong Fezf2 band, and defined the L5 and L6 border as between the strong and weak bands of Fezf2. To account for variation in cortex thickness and sample preparation, we examined three slices spanning 800 µm in the auditory cortex, normalized all cortical thickness to 1200 µm (i.e. the same cortical thickness as the BARseq brains), and calculated the mean positions of layer boundaries (Fig. S3D), The L2/3 and L5 border defined by Fezf2 agreed with that defined by Cux2. Based on these measurements, we defined the L2/3 and L5 border to be at 590 µm and the L5 and L6 border to be at 830 µm. These borders were used for the BARseq analyses when layer identities were involved.

We saw few projection neurons in superficial L2/3 in our dataset. This is partially due to smaller number of neurons labeled near the cortical surface, and partially due to an enrichment of neurons without detectable projections in superficial L2/3 (Fig. S3E). Neurons in superficial L2/3 of the auditory cortex were known to project locally and not contralaterally (Oviedo et al., 2010). Because we did not sample neighboring cortical areas, these locally projecting ITi neurons would have shown as non-projecting neurons in BARseq.

### Comparison of BARseq bulk projection pattern to bulk GFP tracing

For bulk projection comparison to GFP tracing data, we used the bulk GFP tracing data from five brains in the Allen connectivity database (Oh et al., 2014) (experiment 116903230, 100149109, 120491896, 112881858, and 146858006; © 2011 Allen Institute for Brain Science. Allen Mouse Brain Connectivity Atlas. Available from: http://connectivity.brain-map.org/). All five brains had cells labeled in the primary auditory cortex and no labeling in non-auditory area. Several areas we collected only corresponded to part of the areas of the same name in the Allen database, including the somatosensory cortex (restricted to the upper-limb, lower-limb, and the trunk areas), the two visual cortices (restricted to mostly area pm and am in the Allen Mouse Brain Connectivity Atlas; these area labels were different from labels in the Allen Reference Atlas), and the striatum (the rostral striatum and the caudal striatum samples were separated by two brain slices, or 600 µm). The auditory cortex area we collected also included the temporal association area. For both BARseq/MAPseq bulk projections and GFP bulk tracing data, we normalized the strengths of projection to individual areas to total projection strengths in all examined areas for that brain first, and then averaged across brains (five brains for GFP tracing and three brains for BARseq/MAPseq). We then calculated the Pearson correlation coefficient and the associated p value between the GFP tracing and BARseq/MAPseq bulk projection strengths of the corresponding brain areas.

### Population analysis of projections

All population level analyses were carried out in MATLAB.

We calculated the false positive rate *FPR* = *N_OB_* / *N_total_*, where *N_OB_* is the number of neurons with barcodes detected in the OB, and *N_total_* is the total number of neurons. We then calculated the false discovery rate *FDR* = (*FPR* × *n*_0_)/*n_p_*, where *n_p_* indicates the total number of projections detected, and *n*_0_ indicates the total number of possible projections that were not detected.

To estimate the number of projection patterns, we binarized the projections using a threshold set by the maximum number of molecules recovered in the olfactory bulb negative control, and counted unique patterns. Because only one neuron out of 6391 had four molecules in the OB, and four out of 6391 had a single molecule each in the OB, this thresholding likely resulted in a conservative estimate of the unique projection patterns. To estimate the contribution of strong projections, we defined a strong secondary projection as one whose normalized count for a particular barcode was at least 10% of the strongest projection for that barcode.

We then estimated whether each projection pattern could have resulted from missing projections due to false negatives. For each “target” projection pattern, we identified all “parent” projection patterns with one more projection. Neurons with the parent projection patterns might have been misidentified as the target projection pattern of interest if a projection was missed. If the ratio of neurons with the target projection pattern to the ratio of neurons with parent projection patterns was smaller than the false negative rate, then we eliminated the target projection pattern. We repeated this process for all projection patterns, and counted the remaining projection patterns. Using a 10 – 15% false negative rate, 237 – 247 of the original 264 patterns remained.

To test if the estimated number of projection patterns was affected by the sensitivity of BARseq/MAPseq, we filtered neurons with a varying threshold for the primary projections (i.e. the strongest projections; Fig. 4E), which resulted in a reduction in projection patterns. To test if such reduction was caused by smaller sample sizes due to filtering, we randomly subsampled the neurons to the same sample size as that caused by thresholding the primary projection. This subsampling was repeated 100 times to estimate the 95% confidence interval.

To estimate the distribution of the strengths of secondary projections in multi-projection neurons (i.e. neurons with two or more projections), we filtered out neurons whose primary projections were too weak to allow more than one barcode molecules in the secondary projections. For example, when calculating the fraction of neurons with secondary to primary projection ratio of 0.1, we only included neurons whose primary projection was at least 20 barcodes. This would allow at least 2 barcodes to be observed in the secondary projection. We then separated the neurons into 100 bins according to their ratio of strengths between secondary and primary projections, and calculated the probability density function from the cumulative density function. We then fitted with a log normal distribution using gradient descent.

### Hierarchical clustering

All clustering analyses (Fig. S5B) were done using the logarithm of the spike-in-normalized projection strength. We first filtered the projection data using non-negative matrix factorization (NMF)(Lee and Seung, 1999) in Matlab. We approximated the projection pattern *X* of *m* neurons to *n* areas as the product of *Y*, an *m* by *k* matrix containing the loadings of the *k* projection modules for each neuron, and *A*, a *k* by *n* matrix containing the projection pattern of each projection module. Each of the *k* projection modules represents a set of projections that correlate with each other. The projection patterns of individual neurons can thus be approximated by a weighted sum of the projection modules (Fig. S5C). We chose a *k* value that captured most of the variance in the data (Fig. S5D) and resulted in similar classification of neurons in the first two hierarchies (Fig. S5E). This resulted in *k* = 6. We then reconstructed the filtered projection data *X*’ = *Y* * *A*. The filtered projection data *X’* was used for clustering.

During each step of the hierarchical clustering (Fig. S5B), we split each node into two groups using k-means clustering on the squared Euclidean distance of the projection patterns. We then evaluated whether the split was significant using a Matlab implementation of SigClust (available from http://www.unc.edu/~marron/marron_software.html). We kept the new clusters if the split was significant after Bonferroni correction and the sizes of the resulting clusters were larger than 1% of all data points. This procedure was repeated for each new node until no new clusters were found.

This clustering did not separate all different binary projection patterns into their own clusters, suggesting that it was conservative. However, it captured high-level differences in projections that were likely reproducible, and were sufficient for our downstream analyses. Additional levels of clustering would not likely affect downstream analyses since most of our findings did not rely on the leaf clusters.

We also compared our clustering to graph-based clustering using Louvain community detection (Blondel et al., 2008) and spectral clustering (Ng et al., 2002) (Fig. S5F). Louvain community detection identified 2-4 clusters at each hierarchy, and therefore did not fully correspond to the clusters obtained by bifurcation only at any hierarchical level. However, the resulting clusters from both methods, especially high-level nodes, were similar to those obtained using k-means. We chose to base all further analyses on clustering using k-means, because the major classes were better separated than using spectral clustering and the imposed bifurcation was easier to interpret than the clusters produced by Louvain community detection.

We then used a probabilistic approach (Tasic et al., 2016) to assign neurons to these clusters. For each pair of clusters, we trained a random forest classifier on 80% of the data. We then used the classifier to classify the remaining 20% of the data. We repeated this process five times, each time using a mutually exclusive group of 20% of data, so that all data were classified once. We repeated this whole process 10 times for each pair of clusters, so that all data were classified between each pair of clusters 10 times. For each barcode, we then removed all cluster memberships that were scored 0 out of 10 in any one of the pairs involving that cluster. The remaining clusters (i.e. ones that have scored at least 1 out of 10 comparisons in any pairwise comparison) were assigned to the barcode, with the main identity as the cluster with the highest sum of scores across all pairwise comparisons involving that cluster. The majority of neurons (5968/6391, or 93% of all neurons) were uniquely assigned to a single cluster with high probability (>98% probability), but a small set of neurons (421/6391, or 7% of all neurons) were assigned to two clusters at around 50% probability each (Fig. S5G).

Although BARseq has a low false-negative rate (∼10%), such a false-negative rate may accumulate for subclusters with multiple projections, leading to higher rate in misclassification. To examine the extent of such misclassification, we classified neurons probabilistically using 10 out of the 11 projections, thus simulating the effect of a projection being uninformative (Fig. S5H). This analysis revealed similar distribution of neurons with the majority being well-classified and a smaller fraction being ambiguously assigned to two or more clusters. We then considered a neuron ambiguously classified if it was assigned to two or more clusters using all 11 projections, or assigned to the wrong cluster using any of the 10 projections if the resulting cluster was consistent with a dropped projection rather than a false-positive projection. This resulted in 5682/6391 (89%) well-classified neurons and 709/6391 (11%) ambiguous neurons. These estimates represent an upper bound on the number of ambiguous neurons, because it did not take into consideration the actual low false-negative rate of BARseq. Therefore the majority of neurons were unambiguously assigned to a single cluster even when potentially missed projections were taken into consideration.

For Louvain community detection, we used a similarity matrix ***S***, in which each element ***S****ij* was the difference between the Euclidean distance of projections between two neurons and the maximum Euclidean distance of projections between any two neurons in the dataset. Louvain community detection was performed using a MATLAB implementation of the algorithm (https://perso.uclouvain.be/vincent.blondel/research/Community_BGLL_Matlab.zip).

For normalized spectral clustering (Ng et al., 2002), we used the same similarity matrix ***S*** as described above. Spectral clustering was performed using a MATLAB implementation of the algorithm (https://www.mathworks.com/matlabcentral/fileexchange/34412-fast-and-efficient-spectral-clustering).

t-SNE(van der Maaten and Hinton, 2008) was performed using a MATLAB implementation of the standard t-SNE (https://lvdmaaten.github.io/tsne/) using the log projection data as inputs.

All high-level subclusters were found in significant numbers across all three brains (Fig. S6A), with the exception for the striatal projecting neurons in the CT branch (Leaf 25 in Fig. 5C). Layer 6 CT neurons, however, usually do not project to the striatum (Shepherd, 2013). The apparent striatal projections could be caused by contamination by fibers passing through the striatum. Alternatively, these neurons could be PT-l neurons in layer 5 and deep layer 6 whose subcerebral projections were missed due to either weak projections in the tectum or projections to other targets rather than the tectum. This is especially likely since we did not sample many potential target areas for PT-l neurons, such as the brain stem. Our analyses of projection neurons *in situ* support the latter hypothesis. We observed a total of 590 corticothalamic neurons that did not project to the tectum or the striatum, and 505 that did project to the striatum. However, only 10 of the latter group was obtained in the *in situ* sequenced brains, compared to 229 of the former group (Fig. S6A). Therefore, most of the striatum and thalamus projecting neurons were from the conventional MAPseq brain (XC14). Because we only sequenced neurons at the center of each injection site *in situ* (XC9 and XC28), but collected a much larger injection site that may have included neighboring cortical areas in the conventional MAPseq brain (XC14), these neurons were rare in the auditory cortex, and were more likely in neighboring cortical areas, where PT-l neurons could project to other targets. Furthermore, the 10 neurons that projected to the striatum had a laminar profile similar to that of PT-l neurons, but different from those of the layer 6 CT neurons (Fig. 5C). These results suggest that most of the striatum projecting “corticothalamic” neurons were likely PT-l neurons in layer 5 and deep layer 6 in neighboring cortical areas. The lack of such neurons in XC9 and XC28 thus likely reflect differences in the locations of neurons sampled due to the differences between MAPseq and BARseq.

### Normalized entropy for the laminar distribution of a group of neurons

We examined whether clustering resulted in subclusters that were more restrictive in laminar locations. One natural measure of spatial compactness is the standard deviation of the spatial distribution of neurons within a class, but such a measure yields spuriously high values for multimodal distributions. We therefore examined the entropy (normalized to fall between 0 and 1) of the laminar distribution of all nodes and leaves in the clustering, a measure which is insensitive to the shape of the distribution.

To calculate the entropy of the laminar distribution of a group of neurons, we discretized the laminar location of the neurons into 13 bins, each covering 100 µm. We then calculated the entropy of the discrete distribution of laminar locations 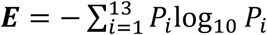, where *P_i_* is the probability of neurons falling into the *i*^th^ bin. We then normalized ***E*** to the maximum possible ***E*** for 13 bins to obtain the normalized entropy ***E***’ = −***E***/log_10_(1/13). The normalized entropy thus equals 0 when all neurons fall into one bin, and 1 when the neurons randomly distribute across all 13 bins.

We did not see significant difference in the entropy of laminar distribution of subclusters across brains (Fig. S6B; p > 0.05 for all subclusters after Bonferroni correction). Furthermore, the distribution of neurons was also consistent across brains: the average laminar locations of neurons from each brain were similar for all but one subcluster (Fig. S6C; p > 0.05 after Bonferroni correction, Mann-Whitney U test). The one exception was CT neurons, which were on average 50 µm deeper in XC28 than in XC9 (Fig. S6C; p < 0.0005 after Bonferroni correction), probably because more deep L6 neurons were labeled in XC28 than in XC9 (XC9 and XC28 had 165 and 132 L6 labeled neurons with laminar depths < 1000 µm, respectively, compared to 56 and 129 L6 neurons with laminar depths > 1000 µm; p < 1e-7 using Fisher’s exact test). Therefore, this observed lack of laminar specificity of subclusters was consistent across samples.

### Single-cell dissociation for RNAseq

To dissociate neurons for single-cell RNAseq, the animal was anesthetized with isofluorane and decapitated. We then dissected out the brain and used a 2 mm biopsy punch to remove the auditory cortex. The auditory cortex was immediately dissected in ice cold HABG medium [40 mL Hibernate A (Brainbits), 0.8 mL B27 (Thermo Fisher Scientific) and 0.1 mL Glutamax (Thermo Fisher Scientific)] into small pieces and placed in 3mL papain solution [3mL Hibernate A-Ca (Brainbits), 6 mg papain (Brainbits), and 7.5 µL Glutamax] pre-warmed to 30 °C in a 15 mL tube. The tube was closed and gently rocked at 30 °C for 40 mins. The digested tissues were then transferred to a tube containing 2 mL HABG pre-warmed to 30 °C and triturated 10 times using a salinized pipette with 500 µm opening. The undissociated tissues were then transferred to a new tube with 2 mL HABG and triturated 10 times using a salinized pipette with 500 µm opening. The remaining undissociated tissues were again transferred to a new tube with 2mL HABG and triturated 5 times. The three tubes of 2 mL HABG each were then combined, and carefully laid on top of a density gradient of 17.3%, 12.4%, 9.9%, and 7.4% (v/v) Optiprep in HABG. This tube was centrifuged at 750 g for 15 mins. The top two fractions were then removed. The next two fractions and half of the bottom fraction, which contained neurons, were collected. The remaining 0.5 mL of the last fraction and the pellet were discarded. The collected neuronal fraction (about 2.5 mL), was diluted in 5mL HABG and centrifuged at 300 g for 5 mins. The pellet was washed in 5 mL HABG, pelleted again, and resuspended in 100 µL HABG. The cell suspension was then kept on ice and processed for library preparation using 10x Genomics Chromium according to the manufacturer’s protocol. The prepared libraries were sequenced using Illumina NextSeq.

### Comparison of barcoded and non-barcoded neurons using single-cell RNAseq

We injected the brain in the auditory cortex as described above. After 24 hours, we dissociated neurons from the auditory cortex as described above, and prepared library using Chromium single cell 3’ reagent kit v2 (10x Genomics). The sequencing library was then sequenced using one lane of Illumina NextSeq. The raw sequencing data were processed using Cell Ranger v2 (10x Genomics) to generate the expression matrix. We then further processed the data using Seurat v2. We first filtered out cells with less than 200 unique genes per cell and more than 10% mitochondria RNA per cell) and log normalized the data. We then regressed out either the effect of the percentage of mitochondria genes linearly (Fig. S2A) or both the effect of the percentage of mitochondria genes and endogenous UMI counts (i.e. excluding barcode reads) using a poisson model (Fig. 3G). We then identified barcoded neurons and non-barcoded neurons by thresholding the level of Snap25 expression and barcode counts. We observed a general reduction in gene expression across most genes, but the relative level of expression among these genes were preserved in barcoded cells (Fig. S2A). Although a minor fraction of genes were over-expressed in barcoded cells, we could further correct for both the under-expression among most genes and over-expression in this subset of genes in the barcoded neurons by simply regressing out the effect of endogenous read counts per cell (Fig. 3G), thus restoring the relative gene expression level to the level of the non-barcoded cells. These results indicate that the relative gene expression level is preserved in barcoded neurons.

To further confirm that the relationship among expression of genes were preserved in individual cells, we examined the amount of variance explained by principal components in neurons (Fig. S2B). We reasoned that if Sindbis infection perturbs major structures in gene expression, then principal components of gene expression in non-barcoded neurons should fail to capture the variance of gene expression in barcoded neurons. We identified genes with variable expression and exported the expression of these genes for both barcoded and non-barcoded cells to MATLAB. We then scaled and centered the data in MATLAB and calculated the explained variance of top principal components. Our results (Fig. S2B) showed that the same top principal components explained similar amount of variance in both barcoded and non-barcoded neurons, thus further indicating that the structures in gene expression is preserved in barcoded neurons. Heatmaps of top genes of principal components (Fig. S2C) were generated in Seurat.

### Single-cell RNAseq of neurons in the auditory cortex

For non-barcoded cells in the auditory cortex, the library was prepared using Chromium single cell 3’ reagent kit v3 (10x Genomics) and sequenced using one lane of Illumina NextSeq. The raw data was processed using Cell Ranger v3 (10x Genomics) followed by Seurat v3 (Stuart et al., 2019). We analyzed 4471 cells with more than 3000 UMI, more than 800 unique RNAs, and less than 15% mitochondria RNA counts. We then linearly regressed out the effect of the percentage of mitochondria RNA counts, and clustered using the top 40 principal components using a resolution of 1 in Seurat. This produced seven clusters (735 cells) that appeared to be neurons with high Snap25 expression. We then pooled these neuronal clusters together and performed a second round of clustering using the top 40 principal components of this subset of data using a resolution of 0.3. This produced the seven neuronal clusters shown in Fig. 6A, B.

We then compared the clusters obtained to those obtained in the visual cortex (Tasic et al., 2018) using MetaNeighbor (Crow et al., 2018). MetaNeighbor was performed on a set of 540 highly variable genes. In this comparison, L4 IT and L5 IT clusters from Tasic et al. (2018) has been merged into a single label “L4/L5 IT”. This analysis revealed that the “CT” cluster in the auditory cortex contained CT neurons, L6b neurons, and seven NP neurons. Furthermore, four PT-l neurons were also co-clustered with IT3 neurons; the small number of PT-l neurons were likely caused by bias during single-cell dissociation and droplet formation. Because our goal was to identify genes differentially expressed across subtypes of IT neurons, we did not attempt to further segregate these neuronal types by clustering, nor did we attempt to enrich for PT-l neurons.

### Correlating projections to transcriptionally defined IT subtypes using BARseq

The four IT subtypes defined by gene expression are largely segregated by laminae (Tasic et al., 2018): IT1 neurons are in layer 2/3; IT 2 neurons are in layer 5; IT3 neurons are in layer 5 and 6; and IT4 neurons are in layer 6. However, because layer 5 contains both IT2 and IT3 neurons, and layer 6 contains both IT3 and IT4 neurons, laminar position alone was not sufficient to distinguish between them. We thus utilized the differential expression of two marker genes, Cdh13 and Slc30a3, to further distinguish these IT subtypes. Cdh13 is highly expressed in IT3, but not IT2 neurons, whereas Slc30a3 is expressed in IT3, but not IT4 neurons.

Based on the gene expression (Fig. 6A) and the laminar distribution of the four IT subtypes, we could distinguish the four IT subtypes using the laminar position of neurons and the expression of Cdh13 and Slc30a3: IT1 neurons are IT neurons in layer 2/3; IT2 neurons are layer 5 IT neurons without Cdh13; IT3 neurons are layer 5 IT neurons with Cdh13, and layer 6 IT neurons with Slc30a3; IT4 neurons are layer 6 IT neurons without Slc30a3. We further included Slc17a7, a highly expressed excitatory neuron marker in the cortex, as a control for both cell morphology and FISH quality.

We collected the same 12 projection targets as in the BARseq-only experiments, but each cortical site was further dissected into superficial and deep layers. This resulted in a total of 18 projection sites. The projection sites were processed as in the BARseq-only experiments and sequenced on a Illumina MiSeq.

The injection site (auditory cortex) was cryo-sectioned to 14 um slices using the home-made tape transfer system. The slides were then fixed and permeated according to RNAscope (ACD Bio) pretreatment procedures for fresh-frozen tissues (cold 4% PFA fixation for 15 mins, followed by dehydration in ethanol and 30 mins of Protease IV treatment at room temperature). We then washed the slide three times in PBST, deactivated residual protease by incubating twice in 40% formamide, 2× SSC, 0.01% Triton-X100 for 10 mins at 60 °C, then washed an additional three times in PBST. We then performed reverse transcription, crosslinking, and neutralization of crosslinkers as in BARseq, except that the reverse transcription was performed for only two hours at 37 °C. We then followed procedures for RNAscope according to the manufacturer’s protocol.

The FISH signals produced by RNAscope were then imaged on a spinning disk confocal using a 20× 0.75 NA Plan-Apo objective with 2× binning. After imaging, the FISH probes were stripped twice in 40% formamide, 2× SSC, 0.01% Triton-X100 for 10 mins at 60 °C. We then continued with BARseq from padlock ligation as described above. Sequencing of barcodes was performed as in BARseq-only experiments, except that the first sequencing cycle was imaged with a 10×0.45 NA Plan-Apo objective and a 20× 0.75NA Plan-Apo objective, both with 2× binning.

The FISH images were then registered to the first cycle high-resolution sequencing images using the DIC channel using intensity-based registration in MATLAB. The high-resolution first-cycle sequencing images were registered to the low-resolution first-cycle sequencing images using intensity-based registration. We then extracted cell-body locations using the “Find Maxima” function in ImageJ, and base-called the barcoded somata from the low-res images. We then filtered out cells with low sequencing quality (below 0.8). We then manually counted FISH reads for excitatory neurons with high FISH quality. Excitatory neurons with high FISH quality is defined by: (1) The neuronal somata were clearly delineated by barcodes; (2) the Slc17a7 reads clearly defined the same soma shape; (3) the neuron did not overlap significantly with neighboring neurons. Cells with fewer than 10 copies of Slc17a7 per cell were further excluded (these might have been cells with low FISH quality, partial cells that were cut during cryo-sectioning, and non-excitatory or non-neuronal cells). The remaining cells, including 781 excitatory neurons from XC75 and 737 excitatory neurons from XC91, were mapped to barcodes at projection sites for further analyses.

### Analysis of projections of IT subtypes defined by gene expression

The projection barcode counts were normalized by spike-ins in the same way as the BARseq-only dataset. The gene expression data were normalized by the mRNA counts of Slc17a7 in each cell.

To assign neurons to clusters defined by the BARseq-only dataset, we first combined barcode counts for the superficial layer and deep layer of each cortical area. We then scaled the barcode count in each area so that the mean projection barcode count in each area is the same in the BARseq/FISH data set compared to the BARseq-only dataset. We then used a random forest classifier trained on the BARseq-only projection data to predict clusters of neurons in the BARseq/FISH dataset.

To quantify the laminar bias of projections to cortical areas, we defined laminar bias as *BC_superficial_*/*BC_superficial_* +*BC_deep_*, where *BC_superficial_* and *BC_deep_* indicates barcode counts in the superficial and deep layers of a cortical area. A bias of 1 thus indicates projections specifically to the superficial layers, and a bias of 0 indicate projections to the deep layers only.

To quantify whether ipsilateral and contralateral projections were more likely to co-occur or were mutually exclusive, we compared the fraction of co-projecting neurons to the fraction of either ipsilateral-only or contralateral-only projection neurons between pairs of IT subtypes defined by gene expression. Among ipsilateral projecting neurons, IT3 neurons were less likely to co-project contralaterally than both IT1 (p = 4e-6) and IT2 neurons (p = 4e-5); IT4 neurons were also less likely to co-project contralaterally than both IT1 (p = 8e-23) and IT2 neurons (p = 4e-17). Among contralaterally projecting neurons, IT3 was more likely to exclude ipsilateral projections than IT1 neurons (p = 0.02). No other significant bias was found among the contralaterally projecting neurons (p = 0.3 comparing IT1 to IT4; p = 1 comparing IT2 to either IT3 or IT4). All statistical significance was obtained using Fisher’s exact test after Bonferroni correction. Therefore, IT3 and IT4 neurons that project ipsilaterally were more likely to exclude contralateral projections than those in IT1 and IT2.

Euclidean distance was used when comparing distances between the projections of IT neurons within or across subtypes defined by gene expression (Fig. 7G). As a control the same analysis was done on IT neurons within or across high-level projection subclusters (Fig. S7E). As expected, most neurons were more similar to neurons within the same projection subcluster compared to neurons in different projection subclusters.

### Retrograde tracing validation of correlation between Cdh13 and contralateral projections

12-week old C57BL/6 animals were injected in the somatosensory cortex with 150 nL CTB-Alexa 488 at 0.6 mm AP, -2.2 mm ML from begma, at 300 and 600 um depths, and in the contralateral auditory cortex with 150 nL CTB-Alexa 647 at 2.3 mm AP, 4.2 mm ML, at 500 and 700 um depths. After 96 hours, the animal was perfused with fresh 4% PFA, post-fixed for 24 hours, and cut to 70 µm coronal sections using a vibratome. Four slices containing the ipsilateral auditory cortex were mounted and imaged for both the green and far red channel on an Ultraview spinning-disk confocal (Perkin Elmer). We then probed for Cdh13 in these slices using RNAscope Fluorescent Multiplex Assay following the manufacturer’s instructions. The Cdh13 probe was visualized with the red Amp4 in the RNAscope kit. We then imaged the green channel, far red channel, and the Cdh13 channel (red), and registered the images to the green/far red channel images taken before RNAscope using those two channels. We then manually counted the Cdh13 expression in ipsilateral projecting or contralateral projecting neurons in layer 6 (defined by a normalized cortical depth of over 830 µm). Rank sum test was used to calculate statistical significance between the distribution of Cdh13 in ipsilaterally projecting neurons and contralaterally projecting neurons.

### Quantification and statistical analysis

All *p* values were produced using the statistical tests as noted in the text. All statistical tests were two-sided. Bonferroni correction was applied to cases involving multiple comparisons, and the corrected p values were reported as indicated in the main text.

### Data availability

All *in vitro* high throughput sequencing datasets (including two BARseq brains, XC9 and XC28, one MAPseq brain, XC14, one brain for the comparison between BARseq and retrograde tracing, XC54, three brains for the combination of BARseq with FISH, XC66, XC75, and XC91, and one brain for BARseq in Cre-labeled neurons, XC92) are deposited to SRA (PRJNA557267 and PRJNA448728). *In situ* sequencing data, images for the MAPseq dissections, and a list of Allen Reference Atlas slices corresponding to the dissection slices will be available from Mendeley Data (doi:10.17632/mk82s9x82t.1, 10.17632/g7kdxznt6w.1, 10.17632/86wf7xfz5x.1, 10.17632/2w649fccnt.1). The same data are provided at https://www.dropbox.com/sh/xz7kwj5aj9yxmee/AAD__4d8DppGA6heaO0EVytua?dl=0 for review purposes.

### Code availability

Analysis code and data needed to reproduce all analyses are available for review purposes at: https://www.dropbox.com/sh/xz7kwj5aj9yxmee/AAD__4d8DppGA6heaO0EVytua?dl=0. The same code and data will be available from Mendeley Data (doi: 10.17632/mk82s9x82t.1, 10.17632/g7kdxznt6w.1, 10.17632/86wf7xfz5x.1, 10.17632/2w649fccnt.1) upon publication.

