## Supplementary material for "High-throughput mapping of long-range neuronal projection using *in situ* sequencing": Fig. S

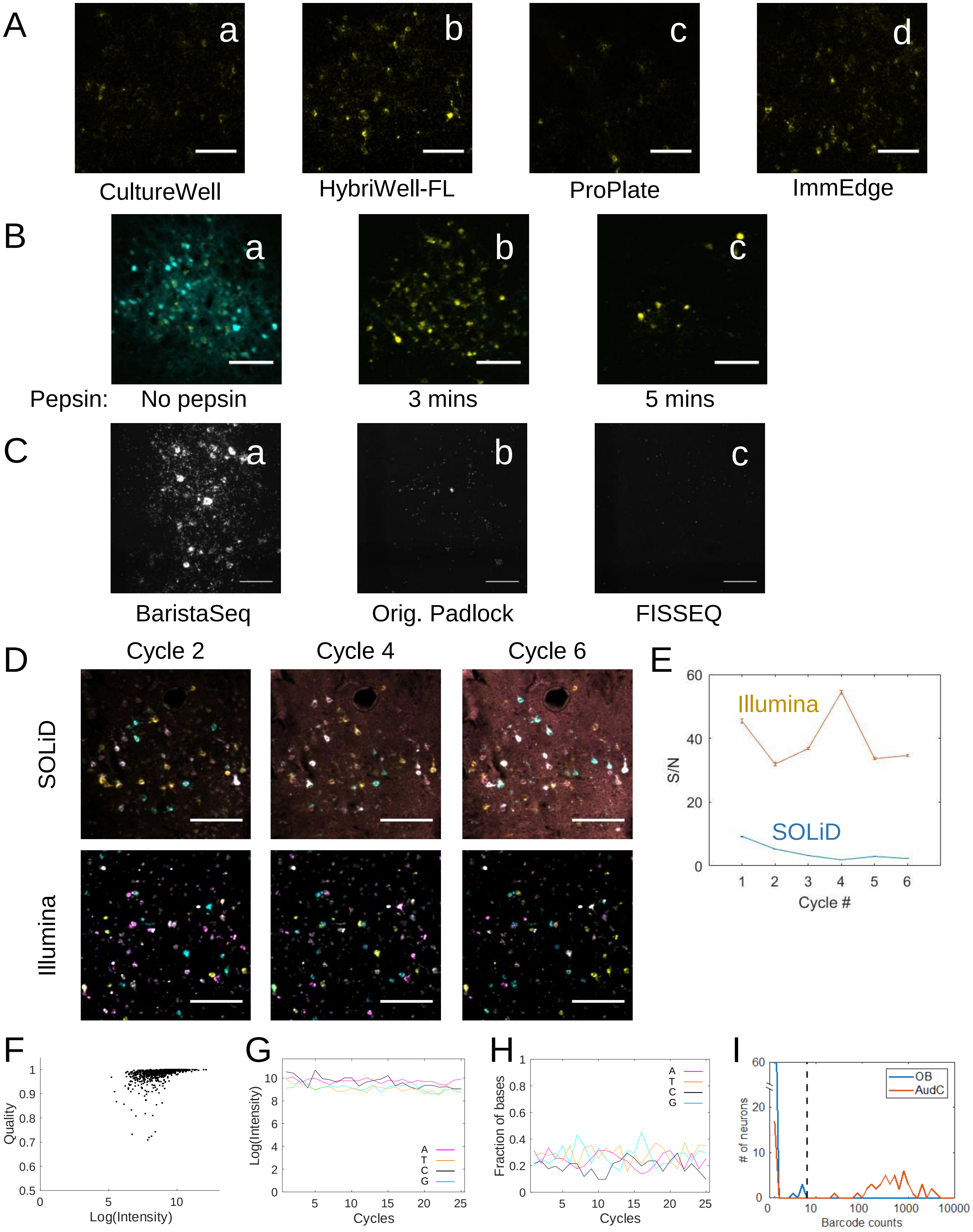


Fig. S1, related to Fig. 2 and STAR Methods

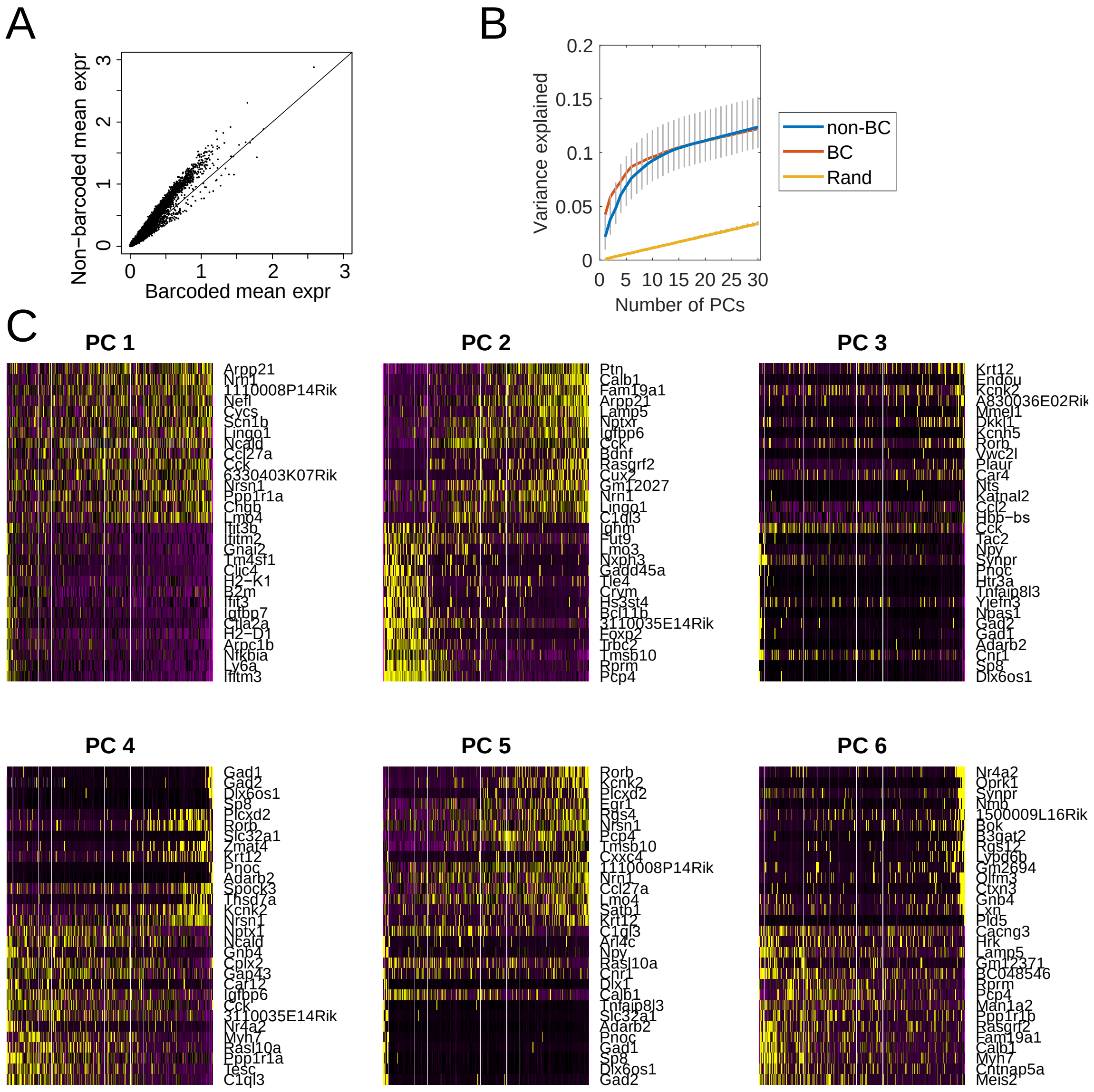


Fig. S2, related to Fig. 3

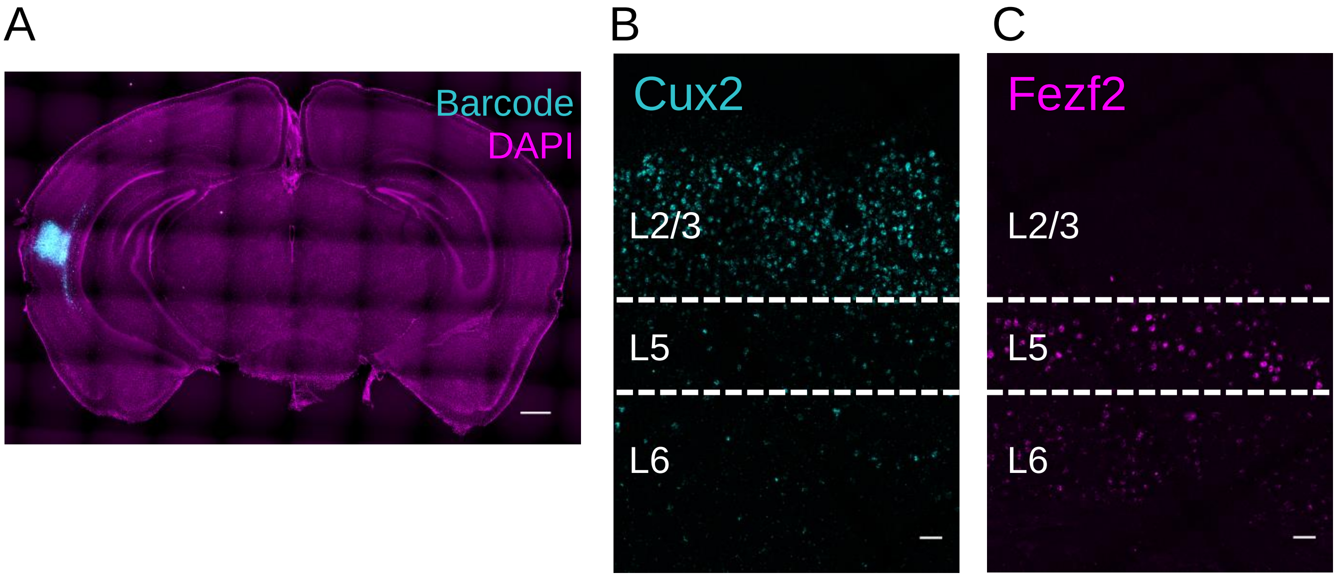


D

| Slice# | 1 | 2 | 3 | Mean ± sem |
|---|---|---|---|---|
| L2/3 – L5 (Cux2) | 640 µm | 590 µm | 550 µm | 600 ± 20 µm |
| L2/3 – L5 (Fezf2) | 550 µm | 600 µm | 590 µm | 580 ± 10 µm |
| L5 – L6 (Fezf2) | 800 µm | 830 µm | 870 µm | 830 ± 20 µm |

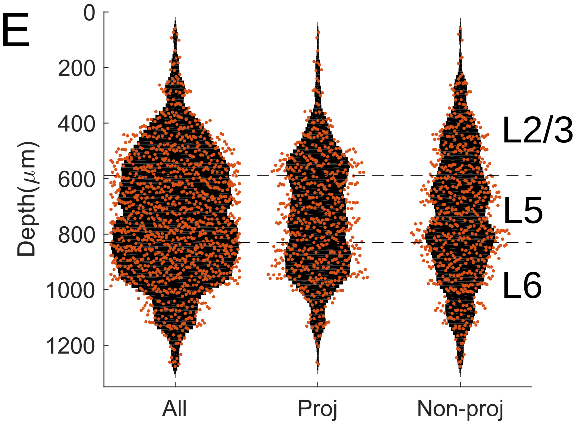


Fig. S3, related to STAR Methods and Fig. 4

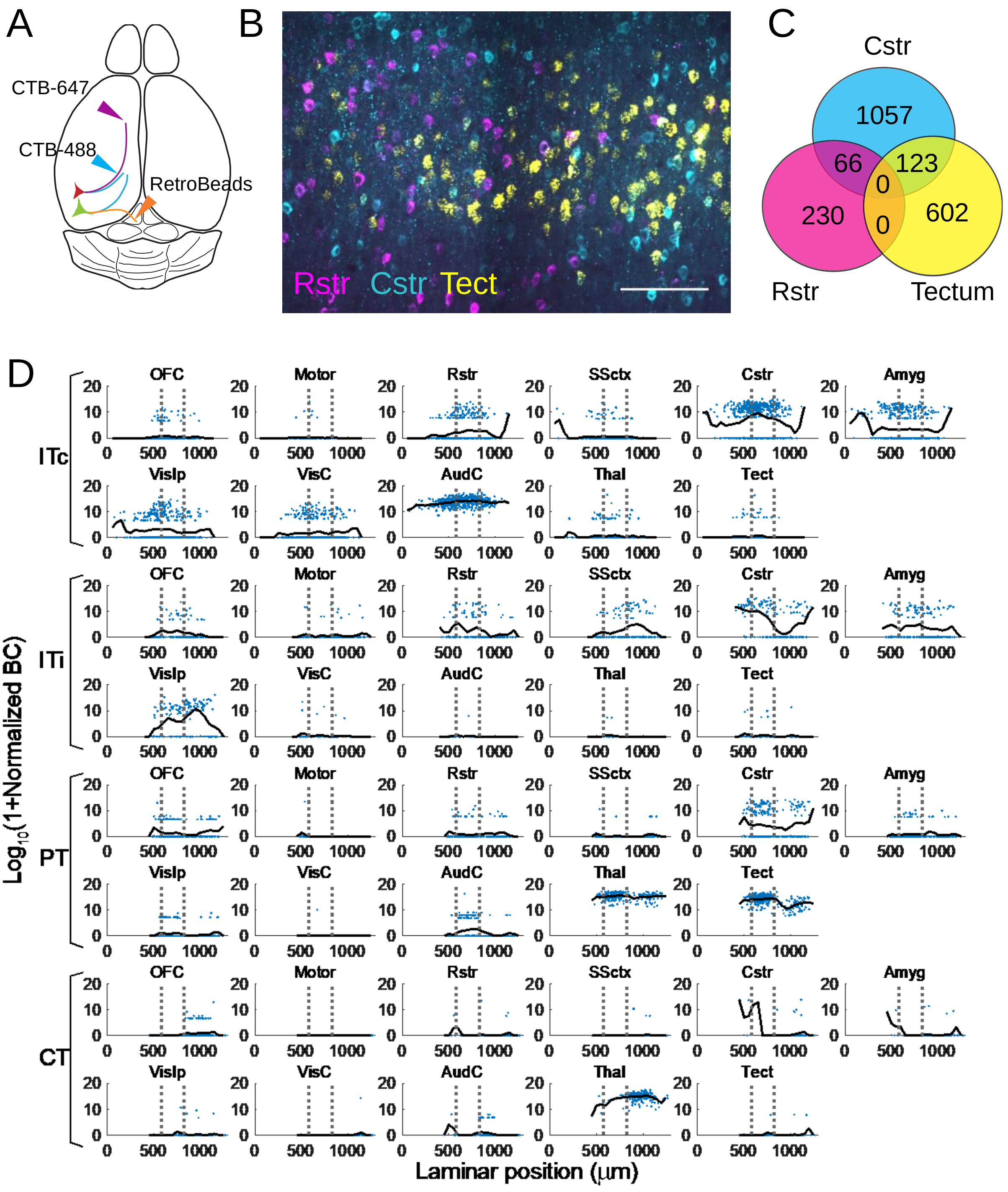


Fig. S4, related to Fig. 5

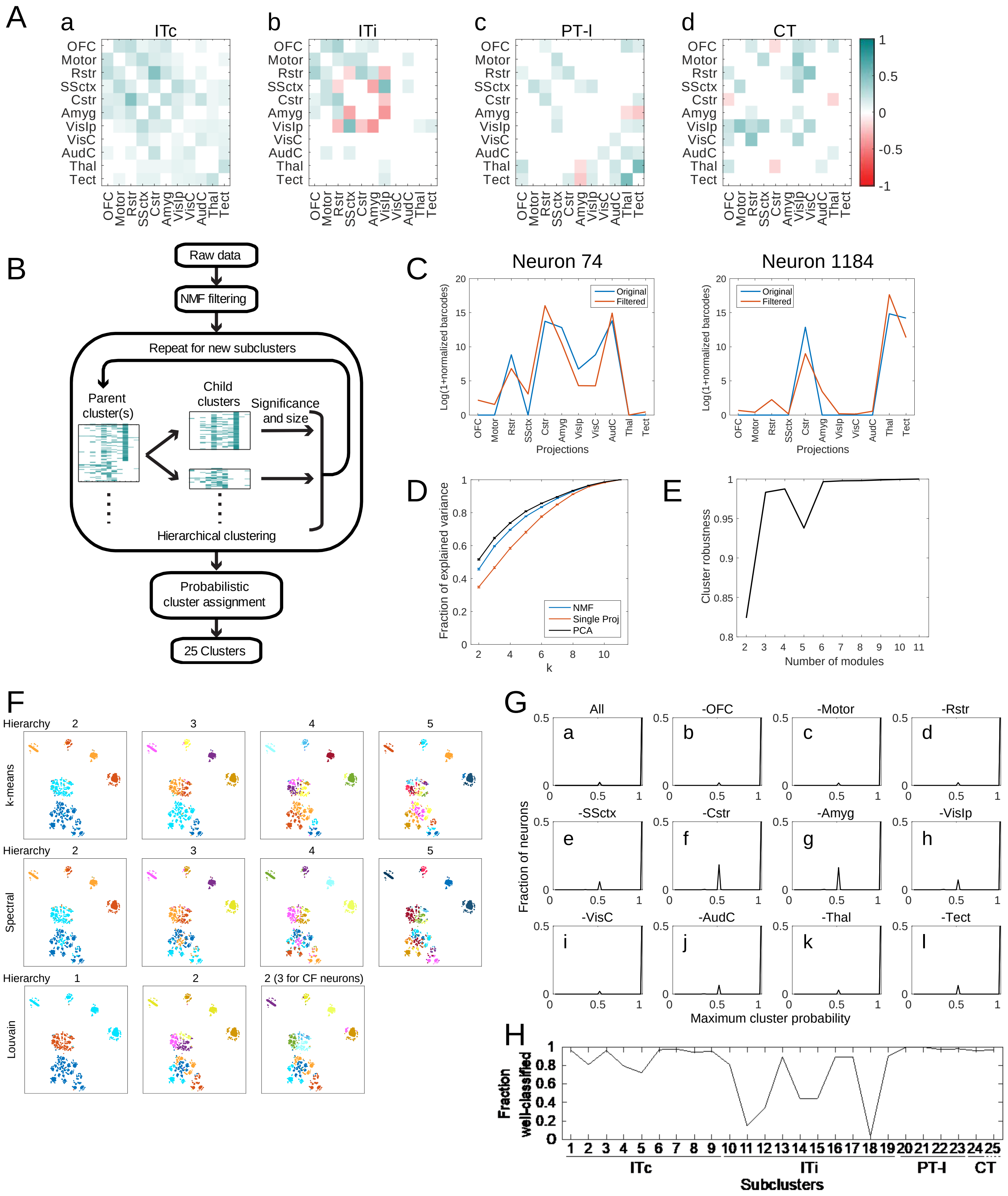


Fig. S5, related to Fig. 5 and STAR Methods

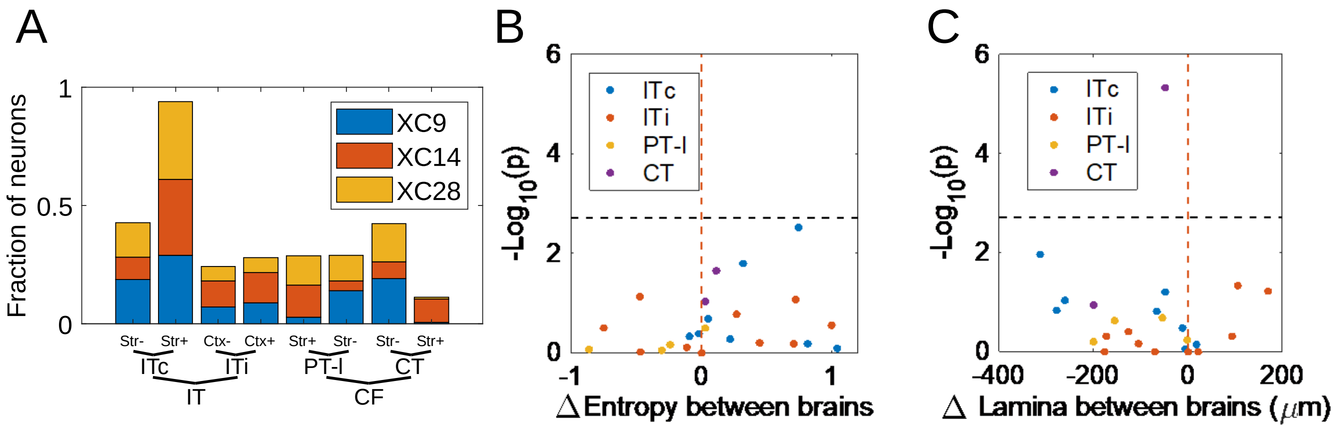


Fig. S6, related to Fig. 5

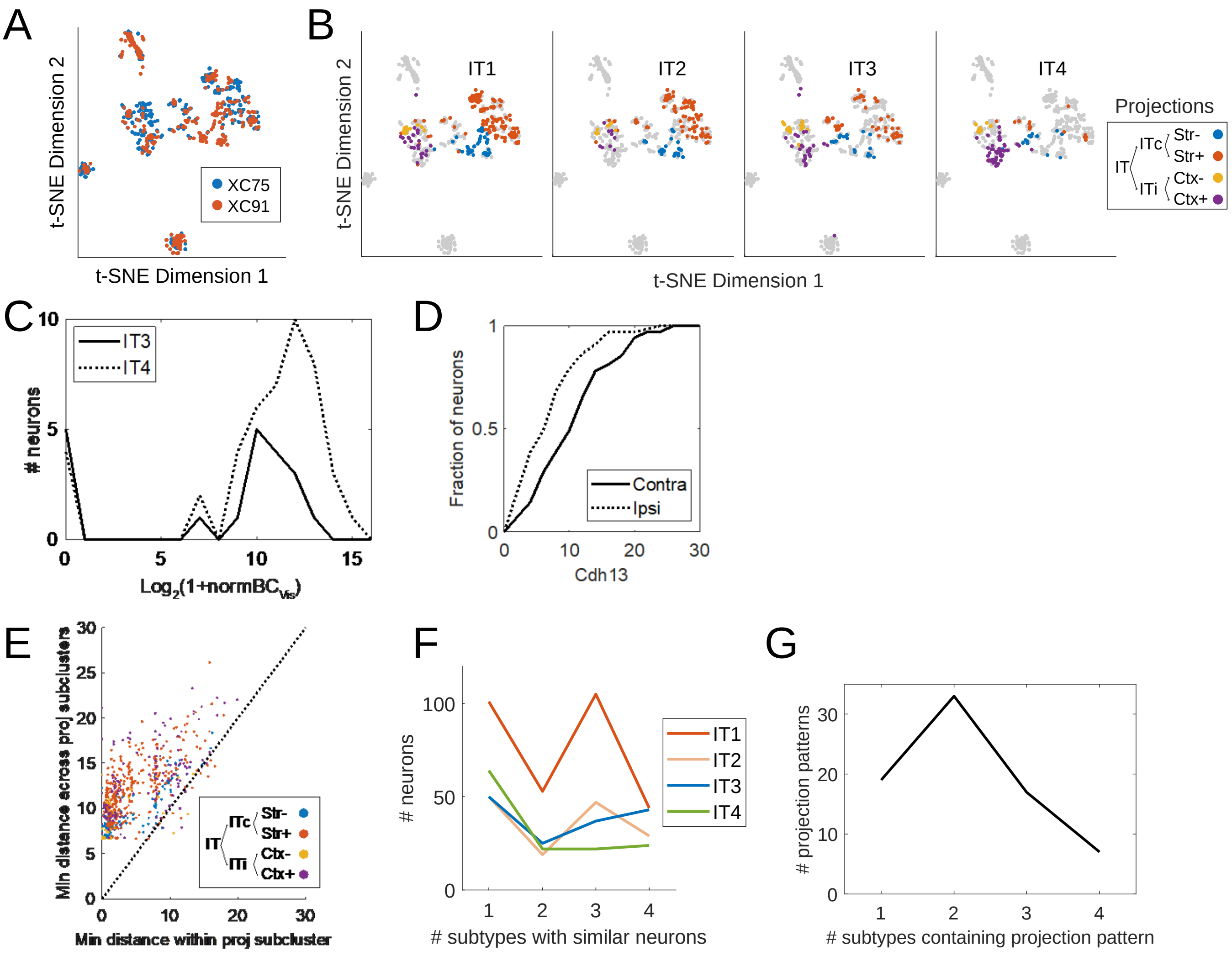


Fig. S7, related to Fig. 6 and Fig. 7

| OB | c1 | c2 | c3 | min qual (>0.75) | CTB+ | BARseq+ |
|---|---|---|---|---|---|---|
| 0 | 151 | 186 | 4 | 0.95 | 0 | 1 |
| 0 | 1433 | 995 | 87 | 0.96 | 0 | 1 |
| 0 | 155 | 70 | 11 | 0.91 | 0 | 1 |
| 0 | 177 | 197 | 17 | 0.89 | 0 | 1 |
| 0 | 17 | 102 | 1 | 0.92 | 1 | 1 |
| 0 | 132 | 135 | 10 | 0.88 | 0 | 1 |
| 0 | 96 | 157 | 12 | 0.88 | 0 | 1 |
| 0 | 102 | 58 | 6 | 0.90 | 0 | 1 |
| 0 | 665 | 58 | 14 | 0.82 | 0 | 1 |
| 0 | 14 | 100 | 5 | 0.87 | 1 | 1 |
| 0 | 0 | 0 | 0 | 0.91 | 0 | 0 |
| 0 | 0 | 0 | 0 | 0.97 | 0 | 0 |
| 0 | 126 | 112 | 7 | 0.84 | 1 | 1 |
| 3 | 281 | 359 | 28 | 0.88 | 0 | 1 |
| 0 | 419 | 313 | 34 | 0.94 | 1 | 1 |
| 0 | 106 | 139 | 3 | 0.79 | 0 | 1 |
| 0 | 72 | 89 | 15 | 0.88 | 1 | 1 |
| 0 | 342 | 159 | 4 | 0.88 | 0 | 1 |
| 2 | 1571 | 1061 | 100 | 0.92 | 0 | 1 |
| 0 | 0 | 0 | 0 | 0.79 | 0 | 0 |
| 0 | 0 | 0 | 0 | 0.86 | 0 | 0 |
| 0 | 0 | 0 | 0 | 0.83 | 0 | 0 |
| 0 | 232 | 51 | 5 | 0.89 | 0 | 1 |
| 0 | 0 | 0 | 0 | 0.92 | 0 | 0 |
| 0 | 372 | 344 | 19 | 0.87 | 1 | 1 |
| 0 | 0 | 0 | 0 | 0.88 | 0 | 0 |
| 0 | 44 | 20 | 1 | 0.83 | 0 | 1 |
| 0 | 234 | 209 | 20 | 0.95 | 1 | 1 |
| 0 | 109 | 10 | 6 | 0.90 | 0 | 1 |
| 3 | 281 | 359 | 28 | 0.91 | 0 | 1 |
| 0 | 61 | 66 | 4 | 0.95 | 0 | 1 |
| 0 | 0 | 0 | 0 | 0.82 | 0 | 0 |
| 0 | 0 | 0 | 0 | 0.95 | 0 | 0 |
| 0 | 335 | 79 | 15 | 0.92 | 0 | 1 |
| 0 | 114 | 186 | 9 | 0.94 | 0 | 1 |
| 0 | 0 | 0 | 0 | 0.90 | 1 | 0 |
| 0 | 271 | 211 | 21 | 0.91 | 0 | 1 |
| 0 | 3159 | 415 | 28 | 0.78 | 1 | 1 |
| 0 | 0 | 0 | 0 | 0.79 | 0 | 0 |
| 0 | 305 | 148 | 18 | 0.94 | 0 | 1 |
| 0 | 0 | 0 | 0 | 0.94 | 0 | 0 |
| 0 | 637 | 293 | 31 | 0.85 | 1 | 1 |
| 0 | 471 | 413 | 15 | 0.93 | 0 | 1 |
| 0 | 1062 | 528 | 94 | 0.92 | 1 | 1 |
| 0 | 335 | 79 | 15 | 0.92 | 0 | 1 |
| 0 | 59 | 73 | 3 | 0.86 | 0 | 1 |
| 0 | 565 | 359 | 27 | 0.93 | 0 | 1 |
| 0 | 460 | 208 | 82 | 0.94 | 0 | 1 |
| 0 | 0 | 0 | 0 | 0.83 | 0 | 0 |
| 0 | 0 | 0 | 0 | 0.84 | 0 | 0 |
| 0 | 0 | 0 | 0 | 0.96 | 0 | 0 |
| 0 | 208 | 69 | 5 | 0.91 | 1 | 1 |
| 0 | 0 | 0 | 0 | 0.91 | 0 | 0 |
| 0 | 1148 | 481 | 30 | 0.97 | 0 | 1 |
| 0 | 200 | 102 | 14 | 0.80 | 1 | 1 |
| 0 | 238 | 154 | 25 | 0.94 | 0 | 1 |
| 3 | 609 | 575 | 62 | 0.88 | 0 | 1 |
| 0 | 18 | 5 | 0 | 0.79 | 0 | 1 |
| 0 | 0 | 0 | 0 | 0.95 | 0 | 0 |
| 0 | 1748 | 163 | 31 | 0.92 | 0 | 1 |
| 0 | 0 | 0 | 0 | 0.96 | 0 | 0 |

### Table S1, related to Fig. 2

| Cell Index | OB | Ipsi-Ctx | Contra-Ctx | Str | Thal | Tect | Total | Slc17a7 | Gad2 |
|---|---|---|---|---|---|---|---|---|---|
| 1012 | 0 | 14 | 87 | 0 | 0 | 0 | 101 | 1 | 0 |
| 1024 | 0 | 119 | 0 | 0 | 0 | 0 | 119 | 1 | 0 |
| 1029 | 0 | 0 | 0 | 0 | 0 | 0 | 0 | 1 | 0 |
| 1074 | 0 | 21 | 18 | 25 | 0 | 0 | 64 | 1 | 0 |
| 1075 | 0 | 0 | 0 | 0 | 0 | 0 | 0 | 0 | 0 |
| 1128 | 0 | 30 | 44 | 10 | 0 | 0 | 84 | 1 | 0 |
| 1135 | 0 | 28 | 3 | 0 | 0 | 0 | 31 | 1 | 0 |
| 1155 | 0 | 0 | 0 | 0 | 0 | 0 | 0 | 0 | 1 |
| 1167 | 0 | 0 | 0 | 0 | 0 | 0 | 0 | 1 | 0 |
| 1179 | 0 | 0 | 170 | 1 | 0 | 0 | 171 | 1 | 0 |
| 1192 | 0 | 0 | 0 | 0 | 0 | 0 | 0 | 0 | 0 |
| 1199 | 0 | 0 | 0 | 0 | 0 | 0 | 0 | 0 | 0 |
| 1226 | 0 | 0 | 0 | 0 | 0 | 0 | 0 | 0 | 0 |
| 1245 | 0 | 0 | 0 | 0 | 0 | 0 | 0 | 1 | 0 |
| 1267 | 0 | 0 | 0 | 0 | 0 | 0 | 0 | 0 | 0 |
| 1274 | 0 | 0 | 16 | 0 | 0 | 0 | 16 | 1 | 0 |
| 2007 | 0 | 4 | 0 | 0 | 0 | 0 | 4 | 1 | 0 |
| 2051 | 0 | 0 | 0 | 0 | 0 | 0 | 0 | 0 | 0 |
| 2054 | 0 | 0 | 0 | 0 | 0 | 1 | 1 | 1 | 0 |
| 3009 | 0 | 0 | 0 | 0 | 0 | 0 | 0 | 0 | 0 |
| 3012 | 0 | 41 | 0 | 0 | 0 | 0 | 41 | 1 | 0 |
| 3076 | 0 | 68 | 48 | 0 | 0 | 0 | 116 | 1 | 0 |
| 3082 | 0 | 9 | 182 | 0 | 0 | 0 | 191 | 1 | 0 |
| 3090 | 0 | 0 | 1 | 0 | 0 | 0 | 1 | 1 | 0 |
| 3130 | 0 | 0 | 74 | 0 | 0 | 0 | 74 | 1 | 0 |
| 4003 | 0 | 34 | 0 | 0 | 0 | 0 | 34 | 1 | 0 |
| 4005 | 0 | 0 | 0 | 0 | 0 | 0 | 0 | 1 | 0 |
| 4008 | 0 | 0 | 0 | 0 | 0 | 0 | 0 | 0 | 0 |
| 4010 | 0 | 253 | 10 | 5 | 0 | 0 | 268 | 1 | 0 |
| 4018 | 0 | 0 | 0 | 0 | 0 | 0 | 0 | 0 | 1 |
| 4022 | 0 | 51 | 51 | 16 | 0 | 0 | 118 | 1 | 0 |
| 4062 | 0 | 0 | 12 | 0 | 0 | 0 | 12 | 1 | 0 |
| 4066 | 0 | 0 | 64 | 1 | 1 | 0 | 66 | 1 | 0 |
| 4067 | 0 | 0 | 0 | 0 | 151 | 0 | 151 | 1 | 0 |
| 4068 | 0 | 11 | 24 | 0 | 0 | 0 | 35 | 1 | 0 |
| 4082 | 0 | 39 | 0 | 0 | 0 | 0 | 39 | 1 | 0 |
| 4088 | 0 | 0 | 0 | 0 | 0 | 0 | 0 | 0 | 0 |
| 4101 | 0 | 10 | 0 | 0 | 0 | 0 | 10 | 1 | 0 |
| 4112 | 0 | 0 | 0 | 0 | 0 | 0 | 0 | 0 | 0 |
| 4114 | 0 | 80 | 57 | 0 | 0 | 0 | 137 | 1 | 0 |
| 4129 | 0 | 0 | 0 | 0 | 0 | 0 | 0 | 1 | 0 |
| 4150 | 0 | 0 | 0 | 0 | 0 | 0 | 0 | 1 | 0 |
| 4163 | 0 | 20 | 4 | 0 | 0 | 0 | 24 | 1 | 0 |
| 4164 | 0 | 0 | 0 | 0 | 0 | 0 | 0 | 1 | 0 |
| 4165 | 0 | 0 | 0 | 0 | 0 | 0 | 0 | 1 | 0 |
| 4169 | 0 | 0 | 0 | 0 | 0 | 0 | 0 | 0 | 0 |
| 4204 | 0 | 0 | 90 | 0 | 0 | 0 | 90 | 1 | 0 |
| 4217 | 0 | 0 | 0 | 0 | 0 | 0 | 0 | 1 | 0 |
| 4224 | 0 | 0 | 47 | 0 | 0 | 0 | 47 | 1 | 0 |
| 4232 | 0 | 2 | 6 | 22 | 0 | 0 | 30 | 1 | 0 |
| 4238 | 0 | 0 | 0 | 0 | 0 | 0 | 0 | 0 | 0 |
| 4241 | 0 | 0 | 105 | 0 | 0 | 0 | 105 | 1 | 0 |
| 4264 | 0 | 14 | 149 | 18 | 0 | 0 | 181 | 1 | 0 |
| 4269 | 0 | 0 | 5 | 0 | 0 | 0 | 5 | 1 | 0 |
| 5001 | 0 | 8 | 106 | 1 | 0 | 0 | 115 | 1 | 0 |
| 5035 | 0 | 4 | 114 | 0 | 0 | 0 | 118 | 1 | 0 |
| 5038 | 0 | 0 | 0 | 0 | 0 | 0 | 0 | 1 | 0 |
| 5076 | 0 | 0 | 80 | 0 | 0 | 0 | 80 | 1 | 0 |
| 5092 | 0 | 0 | 63 | 2 | 0 | 0 | 65 | 1 | 0 |
| 5096 | 0 | 0 | 0 | 0 | 0 | 0 | 0 | 1 | 0 |
| 5111 | 0 | 0 | 0 | 0 | 0 | 0 | 0 | 0 | 0 |
| 5112 | 0 | 0 | 0 | 0 | 0 | 0 | 0 | 0 | 0 |
| 5114 | 0 | 0 | 13 | 0 | 0 | 0 | 13 | 1 | 0 |
| 5144 | 0 | 0 | 3 | 0 | 0 | 0 | 3 | 1 | 0 |
| 5165 | 0 | 7 | 34 | 0 | 0 | 0 | 41 | 1 | 0 |
| 5166 | 0 | 0 | 0 | 0 | 0 | 0 | 0 | 1 | 0 |
| 5178 | 0 | 0 | 0 | 0 | 0 | 0 | 0 | 0 | 1 |
| 6003 | 0 | 0 | 0 | 0 | 0 | 0 | 0 | 1 | 0 |
| 6005 | 0 | 2 | 37 | 1 | 0 | 0 | 40 | 1 | 0 |
| 6030 | 0 | 0 | 22 | 19 | 0 | 0 | 41 | 1 | 0 |
| 6044 | 0 | 42 | 77 | 14 | 1 | 0 | 134 | 1 | 0 |
| 6051 | 0 | 0 | 0 | 0 | 0 | 0 | 0 | 1 | 0 |
| 6077 | 0 | 0 | 0 | 0 | 0 | 0 | 0 | 1 | 0 |
| 6143 | 0 | 0 | 68 | 0 | 0 | 0 | 68 | 1 | 0 |
| 6149 | 0 | 0 | 0 | 0 | 0 | 0 | 0 | 1 | 0 |
| 7006 | 0 | 0 | 0 | 0 | 0 | 0 | 0 | 0 | 0 |
| 7047 | 0 | 0 | 0 | 0 | 0 | 0 | 0 | 1 | 0 |
| 7049 | 0 | 0 | 0 | 0 | 1 | 0 | 1 | 1 | 0 |
| 7063 | 0 | 0 | 0 | 0 | 0 | 0 | 0 | 1 | 0 |
| 7070 | 0 | 0 | 4 | 0 | 0 | 0 | 4 | 1 | 0 |
| 7076 | 0 | 0 | 0 | 0 | 0 | 0 | 0 | 1 | 0 |
| 7083 | 0 | 0 | 0 | 0 | 0 | 0 | 0 | 1 | 0 |
| 7126 | 0 | 0 | 0 | 0 | 0 | 0 | 0 | 1 | 0 |
| 7136 | 0 | 0 | 1 | 0 | 0 | 0 | 1 | 1 | 0 |
| 7140 | 0 | 0 | 24 | 4 | 0 | 0 | 28 | 1 | 0 |
| 8025 | 0 | 36 | 19 | 4 | 0 | 0 | 59 | 1 | 0 |
| 9013 | 0 | 0 | 0 | 0 | 0 | 0 | 0 | 1 | 0 |
| 9024 | 0 | 0 | 0 | 0 | 0 | 0 | 0 | 1 | 0 |
| 9057 | 0 | 116 | 0 | 1 | 2 | 0 | 119 | 1 | 0 |
| 9058 | 0 | 14 | 21 | 0 | 0 | 0 | 35 | 1 | 0 |
| 9063 | 0 | 0 | 0 | 0 | 0 | 0 | 0 | 1 | 0 |
| 9067 | 0 | 12 | 9 | 15 | 0 | 0 | 36 | 1 | 0 |
| 9073 | 0 | 8 | 0 | 0 | 0 | 0 | 8 | 1 | 0 |
| 9078 | 0 | 106 | 0 | 0 | 0 | 0 | 106 | 1 | 0 |
| 9083 | 0 | 0 | 0 | 0 | 0 | 0 | 0 | 0 | 0 |
| 9085 | 0 | 0 | 0 | 0 | 0 | 0 | 0 | 1 | 0 |
| 11012 | 0 | 20 | 10 | 0 | 0 | 0 | 30 | 1 | 0 |
| 11025 | 0 | 0 | 0 | 0 | 0 | 0 | 0 | 1 | 0 |
| 11032 | 0 | 6 | 117 | 0 | 2 | 6 | 131 | 1 | 0 |

Table S2, related to Fig. 3

| | BARseq | | MAPseq* | BARseq + FISH** | | BARseq + Cre*** |
|---|---|---|---|---|---|---|
| Brain ID | XC9 | XC28 | XC14 | XC75 | XC91 | XC92 |
| # targets collected | 12 | 12 | 12 | 18 | 18 | 12 |
| # slices sequenced | 24 | 39 | N/A | 24 | 32 | 10 |
| # of proj barcodes | 4841 | 13581 | 8418 | 16042 | 8388 | 4772 |
| # of cells in ACx | 1575 | 1662 | 13998 | 781 | 737 | 2817 |
| # of cells projecting | 895 | 911 | 5082 | 557 | 422 | 1291 |
| # of filtered cells | 605 | 704 | 5082 | 557 | 422 | 72 (Fezf2+) |

Table S3, related to Fig. 4
